# Overexpression of PSR1 in *Chlamydomonas reinhardtii* induces luxury phosphorus uptake

**DOI:** 10.1101/2022.11.18.517064

**Authors:** Stephen P. Slocombe, Tatiana Zúñiga-Burgos, Lili Chu, Payam Mehrshahi, Matthew P. Davey, Alison G. Smith, Miller Alonso Camargo-Valero, Alison Baker

**Affiliations:** School of Molecular and Cellular Biology, Centre for Plant Sciences and Astbury Centre for Structural Molecular Biology, Faculty of Biological Sciences, University of Leeds, Leeds LS2 9JT, UK; BioResource Systems Research Group, School of Civil Engineering, University of Leeds, Leeds LS2 9JT, UK; Department of Plant Sciences, Downing Street, Cambridge CB2 3EA; Departamento de Ingeniería Química, Universidad Nacional de Colombia, Campus la Nubia, Manizales, Colombia

**Author notes:** Contributed equally to this work (Stephen P. Slocombe, Tatiana Zúñiga-Burgos, Lili Chu). School of Biosciences, Geography and Physics, Faculty of Science and Engineering, University of Swansea, Singleton Park, Sketty, Swansea SA2 8PP (Stephen P. Slocombe, Lili Chu). Scottish Association for Marine Sciences (SAMS) Oban, PA37 1QA, UK.

**Keywords:** Biomass, Micro-algae, Polyphosphate, Transcription, Wastewater remediation, factor

## Abstract

Remediation using micro-algae offers an attractive solution to environmental phosphate (P_i_) pollution. However, for maximum efficiency, pre-conditioning of algae to induce ‘luxury phosphorus (P) uptake’ is needed. Here we show that natural pre-conditioning can be mimicked through over-expression of a single gene, the global regulator PSR1 (Myb transcription factor: Phosphate Starvation Response 1), raising P levels to 8% dry cell weight from 2% in control. Complete removal of P_i_ occurred in log phase, unlike the control. This was associated with increases in PolyP granule size and uptake of Mg^2+^, the principal counterion. Hyper-accumulation of P depended on a feed-forward mechanism, where a small set of ‘Class I’ genes were activated despite abundant external P_i_ levels. This drove a reduction in external P_i_ levels, permitting more genes to be expressed (Class II), leading to more P_i_-uptake. These discoveries enable a bio-circular approach of recycling nutrients from wastewater back to agriculture.

**Teaser:** Manipulating a single gene drove uptake of P and a Mg^2+^ counter-ion for increased PolyP accumulation.

## Introduction

Unlike other macronutrient cycles, the geochemical phosphorus (P) cycle lacks an atmospheric form for replenishment of soils. This has led to an increasing demand from intensive agriculture for mined reserves that are applied as fertilizer. However, inefficient utilization by crops can result in eutrophication of water bodies from agricultural runoff (*1*). Sources of wastewater (i.e. sewage/industrial output) represent a potential source of P, yet sewage treatment plants (STW’s) rarely recycle it back to agriculture or entirely prevent the pollution of waterways (*2*). Circular bioeconomy solutions are needed, and microalgae have a long history in wastewater treatment (*3–5*). Unlike vascular plants, microalgae accumulate P in Polyphosphate granules (PolyP) which can act as a slow-release fertilizer (*6*). This allows algal biomass to be employed in this role, returning P to soils in a controlled manner to minimize runoff (*7–9*). Algae-based solutions have several problems however, including seasonal or climate growth limitations, and there are concerns over the amount of additional land area required for the algal growth. Nevertheless, improvements in P-uptake rates and the P content in biomass could increase the efficiency of the remediation process dramatically (*3, 9*).

Under active growth in P-replete conditions, P resources are assimilated into phospholipids and nucleic acids for cell division (*10, 11*). Producing PolyP would be a diversion from these sinks so presumably cellular processes prevent this happening (*10*) . However, luxury uptake of P can be triggered by restricting other nutrients (e.g. N, S or Zn), resulting in accumulation of PolyP granules in acidocalcisomes (*12*). Alternatively, if P-starvation precedes P-resupply then hyper-accumulation, or ‘P-overplus’ occurs upon reintroduction of inorganic phosphate (Pi), and this is part of the Phosphate Starvation Response (*13*). This response, best understood in *Chlamydomonas reinhardtii*, acts to conserve P, enhance uptake and exploit alternative external P resources (for continued growth) and primes the cell for P hyper-accumulation (*14*). The actual priming phenomenon, which was first recognized in *Micromonas* spp. (*15*), is understood to be a strategic response to fluctuating supplies of P (*13*).

Relying on these natural mechanisms to ensure algal PolyP accumulation in any form of wastewater remediation would require stress pre-conditioning or strict control of nutrients; a process that is already constrained to fit into existing wastewater treatment pipelines (e.g. STW’s) (*4*). An alternative is to exert molecular control over the Phosphate Starvation Response. This includes conservation measures such as replacement of phospholipids with sulfolipids (*16*) and shifts in P_i_ importer gene expression (e.g. repression of *PTA 1,3* and elevation of *PTB 2-5,8*) that result in substantial increases in uptake rate and affinity for P_i_ (*17, 18*). Additionally, there is an induction of periplasmic phosphatase activity (e.g. *PHOX*) to release P_i_ from external organic P_i_ sources (such as glucose-1-P). Finally, there is increased capacity for PolyP synthesis such as the upregulation of *VTC1, VCX1* that encodes respectively a subunit of the transmembrane PolyP synthesis complex and vacuolar Ca-importer (*18*).

The great majority of the above changes are dependent on a single gene: *PSR1* (Phosphate Starvation Response 1), which encodes a Myb-type transcription factor. This was determined by the effects of a non-lethal knockout mutant *psr1-1* (*14, 16–18*) which appears to have a complete loss-of-function: see Suppl. info in (*19*). The *PSR1* gene has global influences, including induction of storage lipid synthesis associated with S-, N- (*19*) and P-deprivation (*16*). These last two studies focused primarily on organic storage products and found that over-expression of PSR1 led to starch increases (and expression of associated genes) along with increased cell size (*16*). High levels of storage lipid in a sub-population of large-celled ‘liporotunds’ was also observed (*19*). Reports of a transient 5-fold increase in P-content per cell were noted in one of the PSR1 over-expression studies (*16*). A complete analysis of the P-uptake and P-storage characteristics, including PolyP content or the gene expression changes promoted by PSR1-over-expression, is absent from the wider literature.

Our aim in this study was to focus on the effect of PSR1-over-expression on P-uptake in *C. reinhardtii*, paying close attention to removal rates and cellular levels of PolyP. We were also interested in relating the dynamics of P-uptake and PolyP synthesis to global patterns of gene-expression. This would reveal how P-homeostasis can be manipulated for harnessing the true potential of microalgae for remediation.

## Results

### PolyP accumulation is increased with PSR1-OE

Growth under replete conditions led to transiently enhanced PolyP accumulation and granule size with PSR1-OE (**Fig. 1**). This was achieved by constitutive PSR1-over expression (PSR1-OE) in *C. reinhardtii* strain UVM4 (*20*). A C-terminal PSR1-YFP fusion (predicted size 1048 amino acids or 109.4 kDa) (see Methods, fig. S1-3, table S1, 2) was expressed in three independently-transformed lines (8-27, 8-2 and 8-42) (fig. S4). Confocal analysis indicated nuclear targeting of the PSR1-YFP fusion in line 8-27 (**Fig. 1B-D**) and line 8-42 (fig. S5B-D) whereas PolyP granules were located to the acidocalcisomes or vacuolar bodies (**Fig. 1E** for 8-27; fig S5E for 8-42). In transgenic line 8-27, PolyP granules were found to be much larger in day 2 of culture (**Fig. 1L**) compared with the UVM4 control (**Fig. 1I**). The signal was diffuse in both lines at this stage however. By day 3, granules had become smaller and more numerous in both lines although the signal was greater in the control (**Fig. 1J** cf. **Fig. 1M**). By day 6 the signal was diffuse and weak in both lines (**Fig. 1K, N**) (multiple images shown in fig. S6).

**Fig. 1.**
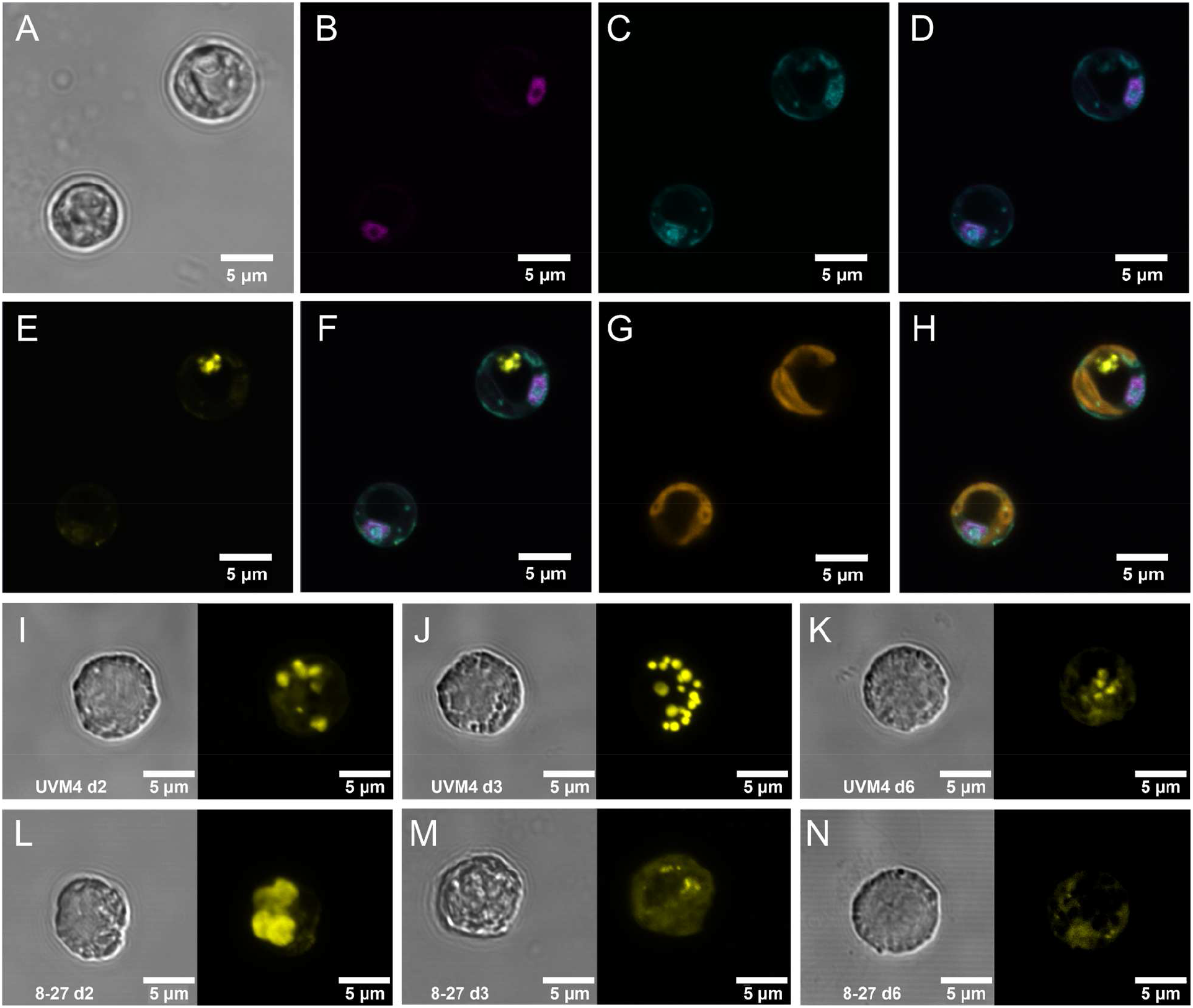
Intracellular localization of PSR1-YFP fusion protein and associated increases in PolyP storage granules. Intracellular localization, as determined by fluorescence confocal microscopy, of the PSR1-YFP fusion protein shown in (**A-H**) for two representative cells from PSR1-OE line 8-27 grown in TAP media. (**A**) bright-field images indicating cell diameter. (**B**) Venus-YFP signal (Emission λ 520-550 nm: magenta) indicating targeting to the nucleus which was identified by DAPI-DNA fluorescence (Emission λ 420-475 nm: cyan) (**C**), followed by co-localization of the DAPI and YFP signals in the merged image (**D**). The PolyPhosphate (PolyP) granules are indicated by the DAPI-PolyP (Emission λ 535-575nm: yellow) (**E**). These are visible as separate entities from the DNA-DAPI stain in the merged image (**F**). Chlorophyll UV-fluorescence (Emission λ 670-720 nm: orange) indicating the single large cup-shaped chloroplast (**G**) and the merged image (**H**) placing the PolyP signal to the periphery of the dark central region of the cell (vacuole). Displayed in (**I-N**) are differences in the accumulation of PolyP granules in cells from a batch culture time course in TAP media (**Fig. 2**) comparing Replicate 1’s of the control line UVM4 (**I-K**) and the PSR1-OE line 8-27 (**L-N**) at three different time points (indicated). Each panel is split between bright field (left) and the DAPI-PolyP signal (right) (Emission λ 535-575nm). A representative cell image was taken from multiple cell images (fig S6).

### Growth rate is unaffected by PSR1 over-expression

No significant changes in growth rate (fig. S7A) or biomass dry weight (DW) concentration (**Fig. 2A**) were noted between control and transgenics. The three PSR1-OE lines (8-2, 8-27, 8-42) were compared with an untransformed UVM4 control in batch culture (25°C under continuous light in TAP medium: 1 mM P or ~30 mg/L P). Exponential growth indicative of log phase was evident from d0 to d2, with no recorded lag-phase, and stationary phase was evident from d4 (fig. S7A). Maximum growth rates were 1.4 d^-1^ (td ~ 0.5 d) with maximum biomass productivity at 0.18-0.19 g DW L^-1^ d^-1^ (**Table 1**).

**Fig. 2.**
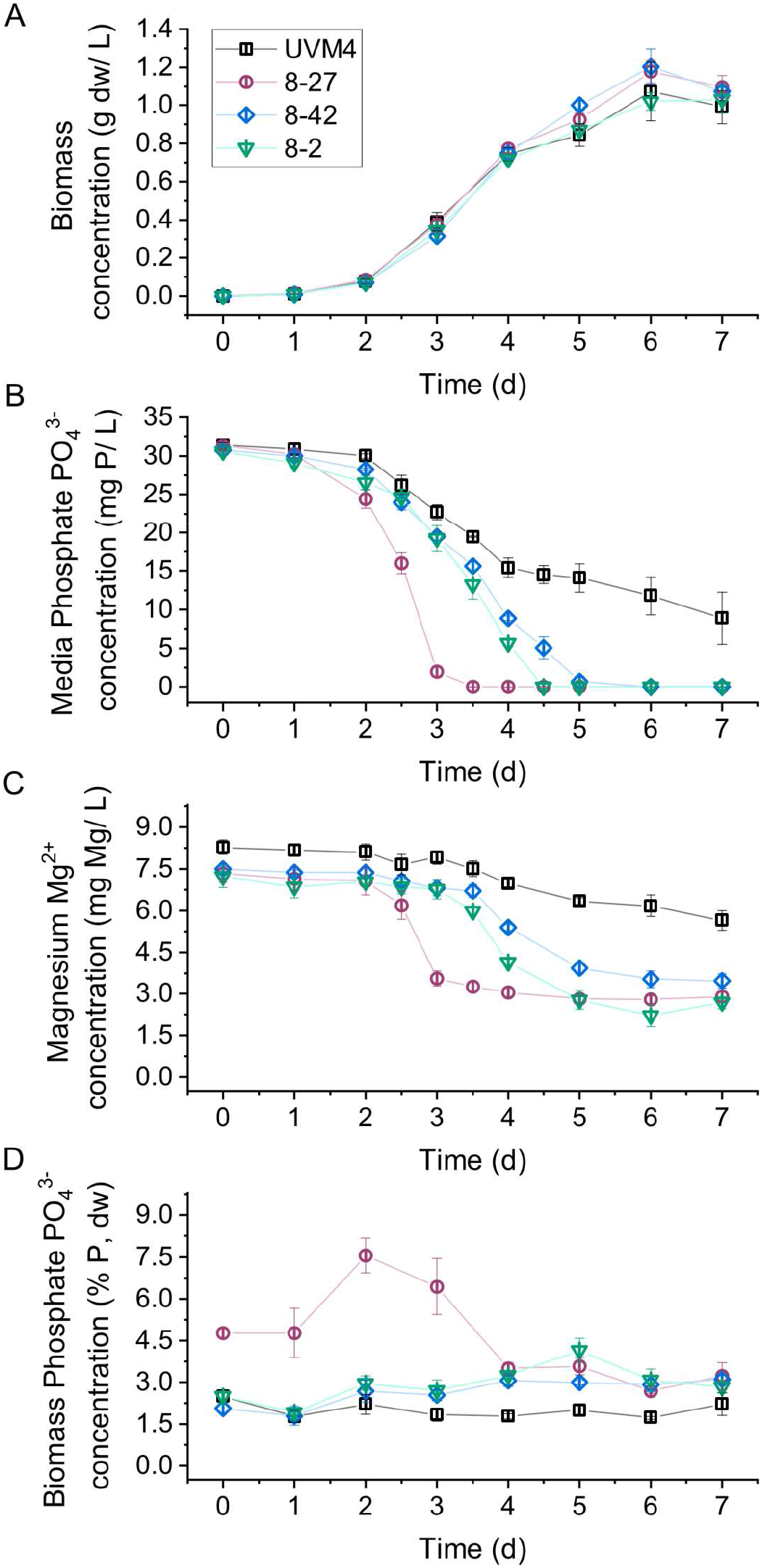
Enhanced phosphate removal and accumulation in PSR1-OE lines. Measurements are shown of post-filtration culture medium composition and biomass composition at different stages of algal growth. Three transgenic PSR1-OE-lines, along with untransformed UVM4 background control, were cultivated under small-scale batch culture conditions in TAP media (30 mg/L P ~ 1mM) in continuous light. The following parameters were monitored over time (**A**) Biomass DW concentration; (**B**) PO_4_^3-^ in media determined by colorimetric assay; (**C**) Mg^2+^ concentration in media determined by ICMS and (**D**) PO_4_^3-^ mass concentration in biomass by assay. Measurements of pH, Chlorophyll, N (ammonium), SO_4_^2-^ and Ca^2+^ are shown in fig. S7.

**Table 1.**
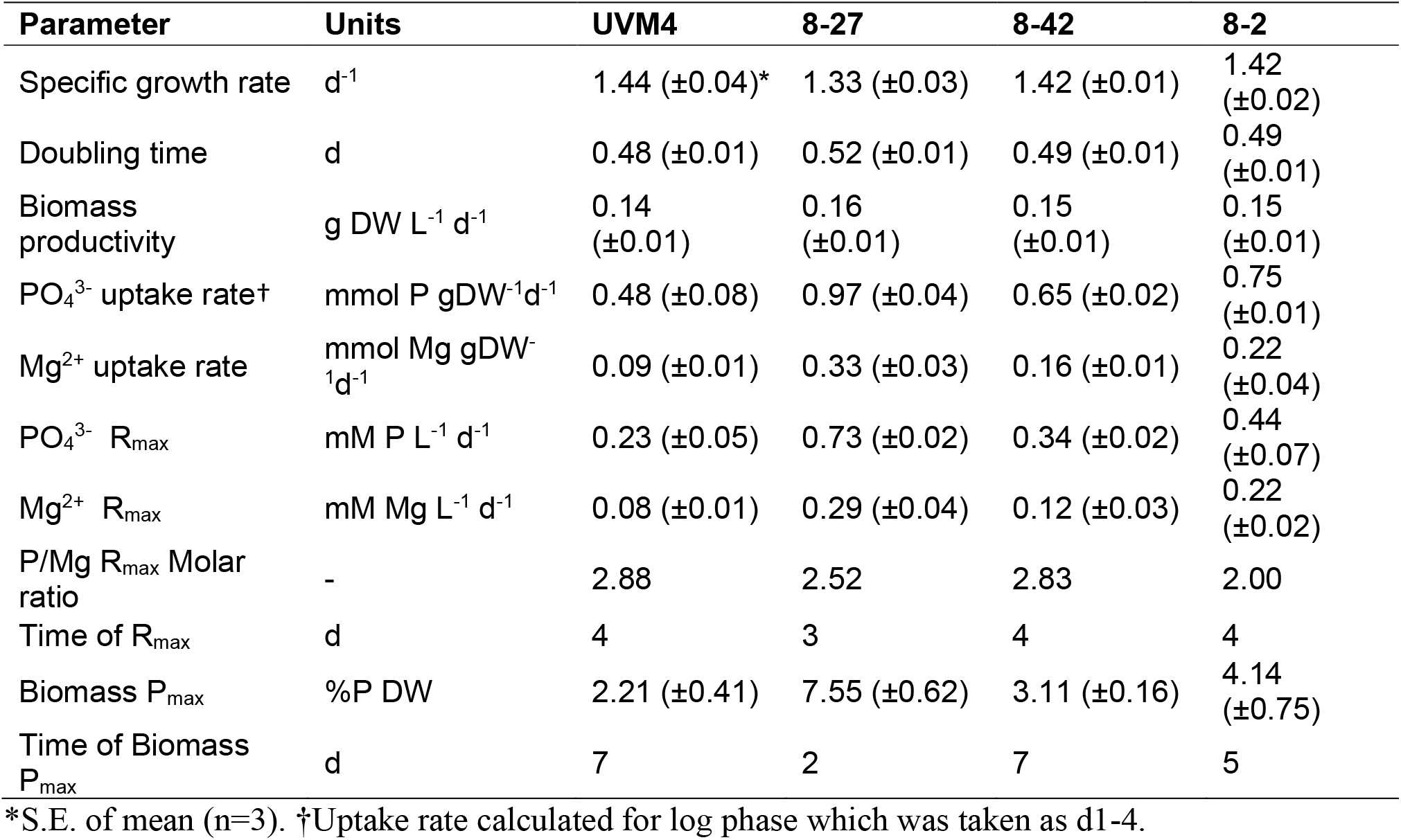
Physiological parameters during PSR1-OE in batch culture over time (7 d)

### Enhanced removal of phosphate from the medium in PSR1-OE lines

Uptake rates for PO_4_^3-^ were substantially improved with PSR1-OE leading to depletion from the medium and uptake of Mg^2+^ was also enhanced (**Fig. 2**, **Table 1**). The removal process was incomplete in the control however.

In UVM4, removal of PO_4_^3-^ and NH_4_^+^ from growth media was down to 30% of starting levels by day 7 (**Fig. 2B** and fig. S7D), SO_4_^2-^ to 50% and Mg^2+^ to 80% (fig. S7E, **Fig. 2C**) and there was little change in K^+^ and Ca^2+^ levels (fig. S7F, G). Parameters that did alter reflected sigmoidal increases in biomass in UVM4 (**Fig. 2A**). This was apparent for pH increases and reductions in PO_4_^3-^, NH_4_^+^, SO_4_^2−^ and Mg^2+^ (**Fig. 2B, C**, fig. S7C-E)

With the exception of PO_4_^3−^ and Mg^2+^, inorganic ions showed no significant change in the transgenic lines against control (**Fig. 2B, C**, fig. S7). Here, the maximum removal rates (R_MAX_) for PO_4_^3−^ and Mg^2+^ were respectively 3- and 4-fold higher for the strongest line (8-27) (**Table 1**). This led to a complete removal of PO_4_^3−^ from the medium by 3.5 d (**Fig. 2B**). Medium Mg^2+^ levels followed PO_4_^3−^ very closely in all the lines except that Mg^2+^ baselined to 5-6 mg/L instead of zero (**Fig. 2C**). In the other PSR1-OE lines (8-2 and 8-42) R_MAX_ (P) was enhanced to a lesser degree (1.9 and 1.5 fold cf. control respectively), and complete removal of PO_4_^3-^from the medium was accomplished later (but not in UVM4) (d4.5 and d5-6 respectively) (**Table 1, Fig. 2B**).

### Hyper-accumulation of biomass P is linked to transience

Rapid P-removal was associated with a greater amplitude and transience of the biomass P peak (**Fig. 2D**). In PSR1-OE line 8-27, the PO_4_^3−^ content in the dried algal biomass reached an early maximum of 8% DW (P) at d2. Line 8-2 peaked later at 4% DW (P) on d5. Line 8-42 reached 3% DW (P) at d4-7 with no distinct peak and the control showed no change at all over time, at 2% DW (P). The higher levels of biomass P in the three transgenic lines were all significantly greater than control levels (Tukey HSD test, fig. S8).

### The Phosphate Starvation Response was weak in the UVM4 control

Under batch culture on P-replete medium (TAP) there is a backdrop of growth limitations and stresses that develop as biomass increases. Gene inductions associated with P-stress were found to be much less evident than for other stresses (**Fig. 3A**).

**Fig. 3.**
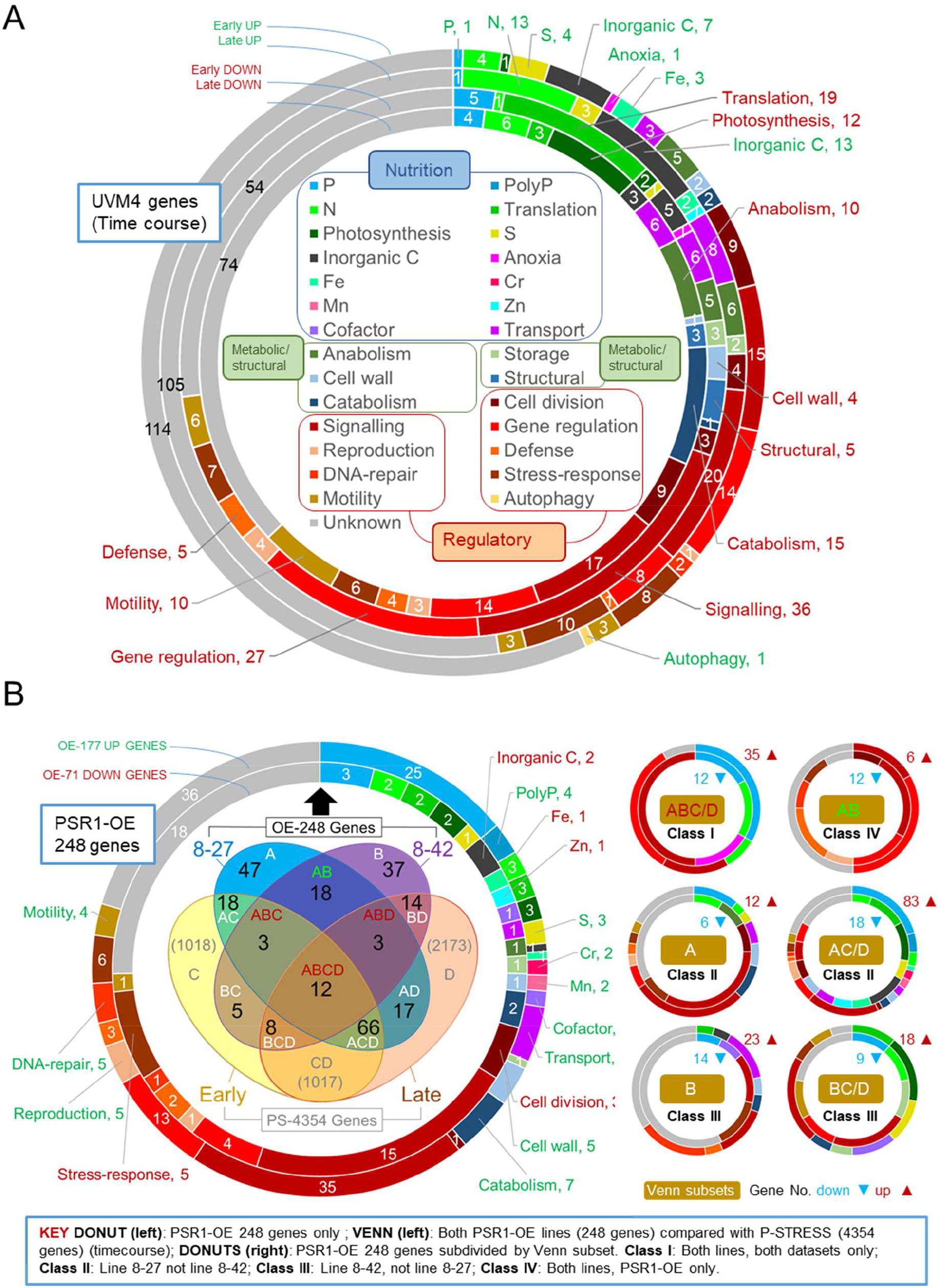
Gene function analysis of transcriptome data. Shown in the upper donut chart (**A**) are relative gene changes occurring within the background strain UVM4 during batch culture, comparing early (d 3 v. d 2) and late (d 6 v. d 2) for up or down changes. Here gene functional roles are shown for the top 200 expression changes in each up/down category (inset). (**B**) Shown an analysis of the FC magnitude expression data for 248 genes (OE-248) having a biologically significant change (>2-fold cutoff) in one or more transgenic lines relative to the UVM4. The large donut chart (left) shows the gene functional analysis for OE-248 for the up (177) or down (71) regulated genes. The Venn diagram (inset) shows a comparison of OE-248 with PS-4354 comprising 4354 genes obtained from the P-STRESS dataset (>2-fold FC cutoff) for “early” (d3) and “late” (d5). Gene number per Venn sector is indicated (in black for OE-248) and each sector is labelled as follows: A (8-27), B (8-42), C (P-STRESS d3) and D (P-STRESS d5). Smaller donut charts (right) shows the gene functional analysis for the six principal Venn sector subsets. These are labelled Class I-IV according to the key (inset). Gene numbers and direction of regulation are indicated.

To examine gene expression, RNA sequencing was performed on time points (d2, 3, 6) from the batch culture experiment in **Fig. 2** for lines 8-27, 8-42 and UVM4 (data S1). In **Fig. 3A**, analysis focused on the UVM4 control as it transitions from log-phase to early stationary phase (top 200 genes; full set in fig. S9 and data S1). Apart from the unknown genes, the largest functional group were regulatory or associated with sexual reproduction, motility or stress responses (30-40%). Genes linked to nutrient limitation or acquisition amounted to 20-30% of the total. Those genes classed as metabolic enzymes or cell structure-related were under 20% of total.

The impact of a particular stress can be estimated from expression changes in associated genes (e.g. gene numbers, magnitudes or early timing). By these criteria, responses to Fe, C, N and S limitation in UVM4 were much greater than those for P **Fig. 3A** (Data S1). Early-induced examples were Fe (FEA1), S (SLT3), or C (CAH8), with mostly late-inductions for N (NIT3, NAR1.2, and GLN3) or P (PTB12) (Data S1). Early gene-induction responses potentially attributable to N/P were AAH1 (amino acid catabolism) and GDP7 (phosphodiesterase, the only early up-regulated P gene) (Data S1). A few (5) down-regulated early response genes were evident for P included PTA3 (P-transport) and a HAD1 (P-hydrolase) (Data S1). Early down-regulation of many protein translation genes (19) were followed by late reductions in photosynthetic (12) and metabolic genes (25) (**Fig. 3A**).

### PSR1 over-expression mimicked P-stress gene induction not repression

The key question was whether the enhanced acquisition of P by PSR1-OE seen under replete batch culture conditions in our experiment was associated with the same gene regulation patterns noted in the literature under P-starvation. We found that in terms of replicating the P-stress response, PSR1-OE was most effective at driving gene induction than repression.

To address this, our data were compared with a published dataset from a P-starvation experiment (P-STRESS) (Data S1) (*16*). The two experiments differed in design where PSR1-OE reflected the evolving changes in the culture during growth whereas in P-STRESS, cells were transferred from P replete to P deficient media at a specific time point. Each approach has advantages and disadvantages, but a comparison was essential to validate the behavior of individual genes or their groups.

Both datasets were time-courses therefore, the fold-change (FC) magnitudes were compared in a Venn diagram with a biological significance cutoff of 2-fold (**Fig. 3B**). This cutoff generated a subset of 248 genes (OE-248) for our data (98% being significant by Bonferroni correction criteria: p-adj<0.1; 68%, p-adj<0.05; Data S1) whereas a subset of ~4k was generated with the P-STRESS data (PS-4354 in **Fig. 3B**) (Data S1). It is important to note that the OE-248 set was a pool of all the significantly affected genes in at least one of the two PSR1-OE lines examined relative to the control.

Good agreement was seen between the PSR1-OE and P-STRESS data sets: 60% of the PSR1-OE set was also altered in the P-STRESS set (**Fig. 3B**). This validation of the OE-248 gene pool was important given differences in the behavior of the two PSR1-OE lines. A higher proportion of genes were co-expressed in the P-STRESS dataset for the strongest line 8-27, (48% cf. 18% in 8-42). There was relatively low agreement between the two transgenic lines at only 15% of the OE-248 genes (**Fig. 3B**). Neither PSR1-OE line showed a co-expression bias towards early- or late-expressed P-STRESS genes (**Fig. 3B**). A bias was seen towards gene upregulation in the OE-248 dataset (71%) compared with downregulation (29%), but not within the P-STRESS dataset (fig. S10 A, B). Upregulated PSR1-OE genes were more likely to follow suit in the P-STRESS data set (55%) than downregulated PSR1-OE genes (27%). These findings also held when a smaller set of highly expressed genes from the P-STRESS dataset were compared to the OE-248 genes, for numerical equivalence (fig. S10 C, D).

### Functional gene categories were regulated differently

Comparing the OE-248 data set with the published P-STRESS data, subdivided the former gene list into six different Venn diagram categories which were condensed into four regulatory classes (I-IV) that differed in gene function profile (**Fig. 3B**).

Regulatory genes dominated in a functional analysis of OE-248 at nearly 50% (either induced or repressed) (**Fig. 3B**, left donut). Nutrient assimilation/partitioning genes accounted for one third of the changes, where half of the induced genes related to P-stress. Unknowns comprised about 25% and metabolic/structural genes had minor representation. For simplicity, each Venn diagram sector was labelled according to the gene groups comprising it i.e. A (line 8-27), B (line 8-42), C (P-STRESS early) and D (P-STRESS late) (**Fig 3B**).

**Class I genes** comprised Venn set ABC/D, a small but robustly substantiated group of 18 genes that were differentially expressed in both transgenic lines (A, B) and the P-STRESS dataset: early (C) or late (D) or both). It was mostly regulatory genes and P-transport genes (e.g. PTB 2-4, PTA1, 3 in **Table 2**) in this grouping (**Fig. 3B**, right donuts). A notable exception was a strongly up-regulated peptidase, GAT1 (**Table 3**).

**Table 2.** Key genes with roles linked to P-homeostasis.

|  |  |  |  |  | Fold-change (FC) |  |  |  |  |  |
| --- | --- | --- | --- | --- | --- | --- | --- | --- | --- | --- |
| Gene | Accession | Class | Description | Correlation | Line 8-27 |  |  | Line 8-42 |  |  |
| P-transport |  |  |  | PSR1 | d2 | d3 | d6 | d2 | d3 | d6 |
| PTB3 | Cre07.g325740 | I (ABCD) | Na<sup>+</sup>/Pi symporter | 0.81* | 26.4 | 26.7 | 8.4 | 4.9 | 2.8 | 3.2 |
| PTB2 | Cre07.g325741 | I (ABCD) | Na<sup>+</sup>/Pi symporter | 0.82 | 8.7 | 11.6 | 9.0 | 2.7 | 2.0 | 1.7 |
| PTB4 | Cre02.g144750 | I (ABCD) | Na<sup>+</sup>/Pi symporter | 0.80 | 9.9 | 7.2 | 8.1 | 2.2 | 1.5 | 1.9 |
| PSTS | Cre01.g044300 | II (ACD) | ABC transporter | 0.83 | 2.8 | 7.9 | 4.6 | 1.3 | 1.2 | 1.3 |
| PTB12 | Cre02.g144650 | II (ACD) | Na<sup>+</sup>/Pi symporter | 0.79 | 1.1 | 7.0 | 3.0 | 0.9 | 0.9 | 0.6 |
| PTB6 | Cre16.g655200 | II (A) | Na<sup>+</sup>/Pi symporter | 0.07 | 4.4 | 0.9 | 0.9 | 1.1 | 0.9 | 1.0 |
| MFT22 | Cre07.g354150 | II (ACD) | Major facilitator permease | 0.96 | 3.0 | 3.9 | 2.7 | 1.5 | 1.2 | 1.7 |
| PTA4 | Cre16.g686850 | II (A) | H<sup>+</sup>/P symporter | 0.28 | 3.7 | 2.1 | 1.2 | 1.1 | 1.1 | 0.86 |
| PTA1 | Cre02.g075050 | I (ABCD) | H<sup>+</sup>/Pi symporter | -0.17 | 2.9 | 0.8 | 0.5 | 2.0 | 1.5 | 0.7 |
| PTB5 | Cre02.g144700 | II (ACD) | Na<sup>+</sup>/Pi symporter | 0.73 | 1.3 | 2.6 | 2.6 | 0.9 | 0.9 | 1.1 |
| PTA3 | Cre16.g686750 | I (ABC) | H<sup>+</sup>/Pi symporter | -0.23 | 2.1 | 1.4 | 1.7 | 1.6 | 1.2 | 0.37 |
| ERM7 | Cre12.g532151 | III (B) | Ca-dependent channel | -0.02 | 0.60 | 0.65 | 0.72 | 0.29 | 0.32 | 0.30 |
| P-salvage/sparing |  |  |  |  |  |  |  |  |  |  |
| PHO5 | Cre04.g216700 | II (ACD) | Exophosphatase | 0.47 | 11.2 | 95.7 | 2.0 | 0.9 | 1.2 | 1.2 |
| GDP2 | Cre16.g683900 | II (ACD) | PLC-like phosphodiesterase | 0.47 | 11.0 | 76.6 | 4.2 | NA | 1.0 | 0.9 |
| MPA2 | Cre09.g404900 | II (ACD) | Metallo phosphatase | 0.50 | 18.6 | 20.7 | 7.7 | 1.6 | 1.2 | 0.9 |
| PHO1 | Cre08.g359300 | II (ACD) | Exophosphatase | 0.54 | 14.7 | 11.9 | 6.1 | 1.2 | 1.3 | 1.3 |
| PWR12 | Cre16.g693819 | II (ACD) | PD-(D/E)XK nuclease | 0.77 | 1.5 | 2.8 | 5.9 | 1.4 | 1.6 | 1.6 |
| EEP2 | Cre06.g295100 | II (AD) | Nuclease/phosphatase | 0.41 | 4.4 | 5.7 | 3.6 | 0.8 | 0.8 | 1.2 |
| Phospho1 | Cre02.g114500 | II (ACD) | Phospholipid phosphatase | 0.93 | 3.7 | 5.2 | 2.7 | 1.7 | 1.3 | 1.6 |
| SPD2† | Cre04.g218200 | II (AC) | Sphingomyelin phosphodiesterase | 0.83 | 1.5 | 4.2 | 3.2 | 0.9 | 1.2 | 0.8 |
| SQD3 | Cre16.g689150 | II (ACD) | Sulpholipid synthase | 0.79 | 1.1 | 2.7 | 3.2 | 1.0 | 1.0 | 1.2 |
| PHT1 | Cre08.g364100 | II (ACD) | Phytase | 0.35 | 1.7 | 3.3 | 0.9 | 0.7 | 1.3 | 0.9 |
| P-homeostasis |  |  |  |  |  |  |  |  |  |  |
| PSR1‡ | Cre12.g495100 | I (ABCD) | Global regulator Myb | 1.00 | 11.6 | 11.1 | 3.3 | 3.1 | 3.0 | 2.7 |
| SPX1† | Cre07.g325950 | II (ACD) | Soluble SPX-protein | 0.89 | 1.7 | 3.2 | 1.8 | 1.4 | 1.4 | 1.3 |
| PolyP synthesis |  |  |  |  |  |  |  |  |  |  |
| PLH1 | Cre02.g078400 | II (ACD) | P-loop NTP hydrolase | 0.56 | 4.6 | 5.7 | 1.4 | 1.5 | 0.8 | 1.4 |
| CAX1 | Cre12.g519500 | II (ACD) | Ca<sup>2+</sup>/H<sup>+</sup> antiporter | 0.73 | 3.2 | 3.1 | 1.9 | 1.5 | 1.3 | 0.7 |
| VTC4 | Cre09.g402775 | II (ACD) | PolyP polymerase catalysis | 0.72 | 1.7 | 2.4 | 1.7 | 1.4 | 1.3 | 1.4 |
| VTC1L | Cre09.g402812 | II (ACD) | PolyP polymerase subunit | 0.90 | 1.7 | 2.4 | 2.1 | 1.2 | 1.3 | 1.5 |
| Cation transport |  |  |  |  |  |  |  |  |  |  |
| MTP4 | Cre03.g160550 | II (ACD) | CDF Mn transporter | 0.53 | 2.5 | 34.9 | 4.5 | 0.8 | 0.7 | 0.7 |
| CIT1 | Cre12.g512950 | II (ACD) | Chromate ion transporter | 0.90 | 2.6 | 5.0 | 2.9 | 1.5 | 1.3 | 1.6 |
| CIT2‡ | Cre12.g507333 | II (ACD) | Chromate ion transporter | 0.89 | 2.7 | 4.8 | 2.2 | 1.6 | 1.2 | 1.2 |
| ACA3 | Cre06.g263950 | II (ACD) | Na<sup>+</sup>/K<sup>+</sup>-exchanging ATPase | 0.82 | 3.4 | 4.6 | 2.7 | 1.6 | 1.6 | 1.2 |
| CPR1 | Cre02.g144550 | II (A) | Na<sup>+</sup>/H<sup>+</sup> exchanger 5-related | 0.77 | 0.9 | 2.8 | 3.0 | 0.8 | 1.1 | 1.47 |
| MTP3‡ | Cre03.g160750 | II (ACD) | CDF Mn transporter | 0.90 | 1.5 | 2.1 | 2.0 | 1.4 | 1.2 | 1.4 |

\*Significant values in bold. Pearson's correlation coefficients (RPKM, p<0.05, n=3). FC values in relation to UVM4 control (p-adj<0.05, n=3). In the *psr1-1* mutant in P-STRESS: †Ectopic down; ‡Independent up (see Data S1).

**Table 3.** Key genes with roles unassigned or linked to processes other than P-stress.

|  |  |  |  |  | Fold-change (FC) |  |  |  |  |  |
| --- | --- | --- | --- | --- | --- | --- | --- | --- | --- | --- |
| Gene | Accession | Class | Description | Correlation | Line 8-27 |  |  | Line 8-42 |  |  |
| Transcription factors |  |  |  | PSR1 | d2 | d3 | d6 | d2 | d3 | d6 |
| SPL9 | Cre16.g683953 | I (ABCD) | Squamosa promoter binding | **0.89\*** | **11.8** | **26.3** | **25.2** | 1.6 | 1.8 | **6.1** |
| Bzip1 | Cre16.g671200 | I (ABCD) | Basic-leucine zipper (bZIP) | **0.96** | **5.5** | **13.8** | **4.9** | 1.3 | 1.9 | **2.8** |
| ZAF1\|\| | Cre26.g756897 | III (BD) | PHD-type C-terminal zinc finger | 0.29 | 0.9 | 1.3 | 0.8 | **2.7** | **4.2** | **10.9** |
| NOR1 | Cre07.g331401 | II (ACD) | Orphan nuclear receptor | **0.85** | **2.5** | **9.2** | **5.2** | 1.1 | 1.3 | 1.4 |
| CSB57 | Cre14.g618820 | IV (AB) | Transposase DNA-binding | **0.69** | **5.3** | **7.3** | **5.9** | **5.0** | 5.5 | 7.0 |
| NOR2 | Cre15.g637552 | IV (AB) | Orphan nuclear receptor | 0.20 | **4.5** | 1.8 | 0.9 | **2.6** | 1.5 | 1.0 |
| RHC1 | Cre02.g144802 | II (A) | RNA helicase | **0.67** | 1.8 | **4.2** | 2.2 | 1.0 | 1.1 | 1.1 |
| ZF3§ | Cre16.g681100 | II (ACD) | Zinc/RING finger C3HC4 domain | **0.92** | **2.4** | **4.1** | **2.5** | 1.2 | 1.3 | 1.5 |
| ZF4 | Cre19.g751047 | IV (ABD) | CCHC-type zinc finger | -0.04 | **3.3** | 1.6 | 2.2 | **2.8** | 1.7 | 1.9 |
| ZF1 | Cre17.g742800 | IV (AB) | CCHC-type C-terminal zinc finger | -0.43 | **0.19** | **0.17** | **0.09** | **0.21** | **0.21** | **0.13** |
| Signal transduction |  |  |  |  |  |  |  |  |  |  |
| MKK5 | Cre10.g463500 | II (ACD) | Mitogen-activated PKK | **0.79** | **4.9** | **26.0** | **3.9** | 1.2 | 0.9 | 0.6 |
| ANK1 | Cre01.g019550 | II (ACD) | Ankryin repeat protein | **0.88** | **5.3** | **14.1** | **6.1** | 1.5 | 1.3 | 1.3 |
| MRK1 | Cre16.g674900 | I (ABCD) | Ser/thr protein kinase | **0.84** | **5.3** | **13.0** | **11.9** | 1.2 | 1.4 | **3.8** |
| AUR1 | Cre16.g674065 | II (ACD) | Aurora protein kinase | **0.75** | **6.6** | **7.1** | **5.2** | 1.2 | 1.1 | 0.7 |
| MEKK | Cre08.g368600 | I (AB) | MEKK-related protein kinase | **0.90** | **6.0** | **6.4** | **5.7** | **2.2** | 2.4 | 1.8 |
| CDPK | Cre02.g114750 | III (BCD) | Ca/calmodulin-dependent PK | 0.19 | 1.3 | 1.4 | 0.8 | **3.3** | **3.7** | **4.1** |
| XBAT31† | Cre09.g403500 | I (ABCD) | E3 ubiquitin ligase | **-0.87** | 0.6 | **0.14** | **0.06** | 0.9 | 0.8 | **0.28** |
| PK7 | Cre08.g369667 | IV (AB) | Thr-specific protein kinase | -0.42 | 0.6 | **0.28** | **0.34** | 0.6 | **0.28** | **0.35** |
| LAR1 | Cre12.g541550 | I (ABCD) | Las17-binding actin regulator | 0.61 | **14.3** | **38.4** | **8.8** | **2.9** | 2.1 | 1.9 |
| BAR | Cre06.g299500 | II (A) | Endocytosis regulation | -0.43 | **0.24** | **0.15** | 0.44 | 1.6 | 1.4 | 1.4 |
| S-stress |  |  |  |  |  |  |  |  |  |  |
| SUTA† | Cre02.g095151 | III (BD) | Sulfate-transporting ABC-2 type | 0.47 | 0.7 | 1.1 | 1.4 | 0.7 | 0.9 | **2.6** |
| SIR† | Cre16.g693202 | III (BD) | Sulfite reductase (ferredoxin) | 0.45 | 0.50 | 1.3 | 1.4 | 0.8 | 1.5 | **2.5** |
| ATS1† | Cre03.g203850 | II (ACD) | ATP-sulfurylase | 0.27 | 0.8 | 0.5 | **0.40** | 0.6 | 0.7 | 1.7 |
| N-stress |  |  |  |  |  |  |  |  |  |  |
| GAT1† | Cre01.g004900 | IV (ABCD) | Glutamine amidotransferase | **0.69** | **5.2** | **3.7** | 2.2 | **4.4** | **2.5** | 1.8 |
| NSI1 | Cre11.g476026 | II (A) | N starvation induced COV1 | **0.80** | 1.3 | **3.0** | **3.3** | 1.0 | 1.1 | 1.3 |
| Other stress responses |  |  |  |  |  |  |  |  |  |  |
| MGS1 | Cre07.g331700 | II (ACD) | Minus gamete specific, secretory | **0.69** | **3.0** | **6.6** | 2.0 | 1.4 | 1.1 | 0.7 |
| UVE1 | Cre12.g505100 | III (B) | UV-damage endonuclease | 0.56 | 0.9 | 1.5 | 1.4 | 1.1 | 0.8 | **2.7** |
| NOS1 | Cre16.g683550 | II (AC) | NO synthase ferredoxin reductase | 0.44 | 0.7 | 1.7 | **2.5** | 0.7 | 1.6 | 1.7 |
| FEA1\|\| | Cre12.g546550 | II (AD) | Periplasmic Fe-binding | 0.08 | **2.2** | 1.4 | 1.3 | 1.9 | 1.5 | 1.6 |
| LCI12 | Cre04.g217962 | III (B) | Zinc finger, CCHC-type | 0.52 | 1.4 | 0.8 | 1.4 | **2.2** | 1.7 | 2.6 |
| DNA2 | Cre12.g528200 | IV (AB) | ATP-dependent helicase (DNA) | -0.44 | 0.6 | 0.7 | **0.24** | 0.5 | 0.6 | **0.32** |
| METE† | Cre03.g180750 | III (BCD) | Vit B12-independent met synthase | 0.43 | 0.54 | 1.2 | 1.5 | 1.8 | 1.2 | **2.1** |
| PBP1 | Cre08.g362600 | IV (AB) | Beta-lactamase | -0.33 | **0.49** | 0.7 | **0.32** | **0.38** | 0.5 | **0.16** |
| CRD1 | Cre12.g537250 | II (ACD) | Cyclase in Tetrapyrrole pathway | 0.29 | 0.7 | 1.8 | **0.31** | 1.5 | 1.3 | 1.8 |
| THB2 | Cre14.g615350 | I (ABD) | Thylakoid truncated hemoglobin | -0.31 | 0.8 | 0.6 | **0.28** | 1.2 | 0.7 | 0.5 |
| GST8‡ | Cre11.g467690 | II (A) | Glutathione S-transferase | -0.26 | **0.08** | **0.04** | **0.04** | 0.8 | 1.5 | 0.9 |
| HMG1 | Cre06.g299550 | II (AB) | HMG-CoA reductase | -0.59 | **0.09** | **0.04** | **0.15** | 1.7 | 1.4 | **2.5** |
| LTI1 | Cre08.g368650 | IV (AB) | Lysin-induced mt- secretory | -0.55 | **0.03** | **0.02** | **0.05** | **0.03** | **0.04** | **0.04** |

\*Significant values in bold. Pearson's correlation coefficients (RPKM, $p < 0.05$ , $n = 3$ ). FC values in relation to UVM4 control ( $p\text{-adj} < 0.05$ , $n = 3$ ). In the *psr1-1* mutant, P-STRESS: Ectopic up†/down‡; Independent up§/down|| (Data S1).

**Class II genes** were only altered in the strongest PSR1-OE line (8-27) and comprised the two largest Venn subsets: set AC/D (also present in the P-STRESS dataset) and set A (specific to OE-248) (**Fig. 3B**). There was a similar split between regulation and nutrition as noted in Class I but also more diverse gene functions. P-stress/assimilation genes were a substantial category among the induced genes in this class but the majority were P-salvage or P-scavenging (e.g. PHO5/X, SQD3 and GPD2) (**Table 2**). Relatively few potential P-transporters were present (i.e. PSTS, PTB12) but significantly, there were four genes linked to PolyP synthesis (**Table 2**).

**Class III genes** were only altered in expression in the weaker OE line 8-42 (Venn subset B and BC/D). A slightly smaller group of functionally diverse genes that included two induced S-assimilation genes (SIR and SUTA) and two repressed P-uptake genes (ERM1, 7) (**Fig. 3B**) (**Table 3**).

**Class IV genes** (Venn set AB) comprised a small number of genes that were affected in both PSR1-OE lines but absent from the published P-STRESS set (and therefore novel) (**Fig. 3B**). They consisted entirely of regulatory pathway genes and unknowns, with a bias towards early gene repression rather than induction (**Table 3**).

### Positive autoregulation mechanism revealed for PSR1

The strongest P-uptake response (line 8-27) appeared to be driven by an early peak in activity of transgene PSR1 mRNA (i.e. late log-phase: d 2-3). In both PSR1-OE lines there was an increase in the endogenous PSR1 gene activity along with increases in P-uptake.

The relationship between the PSR1 protein and mRNA levels are shown in **Fig. 4A, B** with reference to physiological data from **Fig. 2**. PSR1 mRNA levels are shown with a breakdown of the wild type-specific (5’UTR or 3’UTR) and transgene-specific (YFP) fragments alongside the complete PSR1 gene mRNA (5 ’UTR, CDS and 3 ’UTR) which was a combination of wild-type and transgene signals (**Fig. 4B**). The strength of the P-uptake response across the lines was in proportion to PSR1 mRNA and PSR1-YFP fusion protein levels (**Fig. 4A, B**). Activation of endogenous wild-type PSR1 was ~10-fold at d3 for line 8-27 (3’UTR or 5’UTR PSR1) in relation to UVM4 (**Fig. 4B**).

**Fig. 4.**
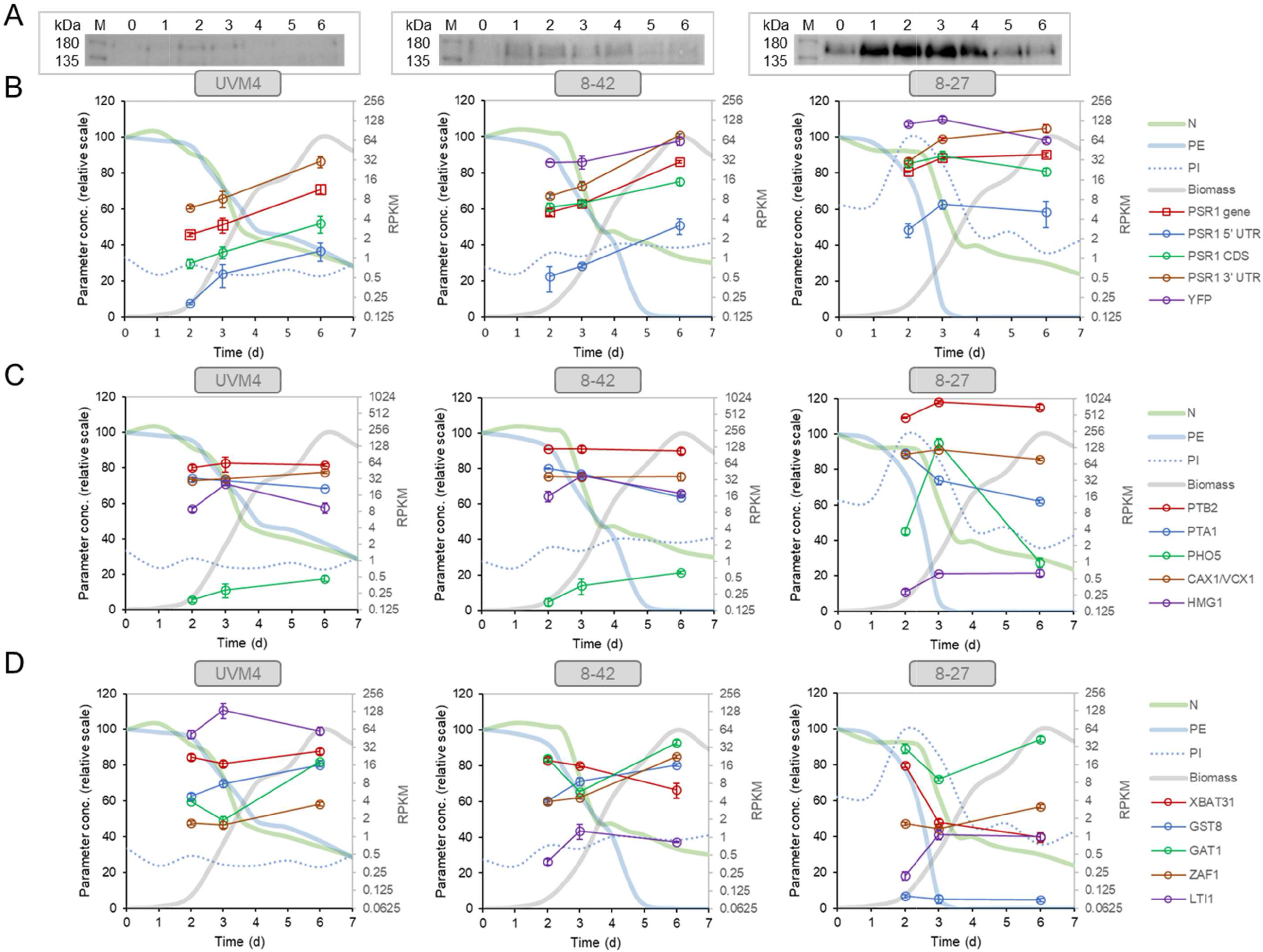
Expression levels of PSR1 and exemplar gene targets elicited by PSR1-OE. (**A**) Inset panels display the western blot signal for anti-YFP antibody showing bands corresponding to the PSR1-YFP fusion protein. Equal chlorophyll loadings (10 μg) were loaded for the batch culture time course, where d0 refers to the starter culture prior to dilution. Gene expression levels determined by RPKM counts are compared over time in batch culture for the transgenic lines (8-27, 8-42) and the background control (UVM4). These data are compared to the physiological data (**Fig. 2**) normalized as follows: N, PE to d0 levels; biomass concentration to maximum in algal line; PI to maximum in experiment. PSR1 mRNA expression is shown (**B**) for the full PSR1 gene sequence and PSR1 CDS (both having combined wild-type and transgene inputs); wild-type-specific fragments (5’UTR and 3’UTR) and construct-specific fragments (YFP). (**C-D**) Exemplar PSR1-OE affected genes are shown illustrating the different classes (Class I-IV) and direction (up/down) of gene regulation: (**C**) Class I-induced (PTB2, PTA1), Class II-induced (PHO5, CAX1/VCX1), Class II-irregular (HMG1); (**D**) Class I-repressed (XBAT31), Class II-repressed (GST8), Class III-induced (ZAF1), Class IV-induced (GAT1) and Class IV-repressed (LTI1). Note all exemplar genes were also affected in the P-STRESS data set (>2-fold change) (*16*). See **Table 2** and **3** for the FC data; full RPKM data in Data S1. Error bars indicate SE, n=3 culture replicates.

The PSR1-YFP fusion protein levels peaked in both transgenic lines at d2-3, although this was only pronounced in line 8-27 (**Fig. 4A**). Total PSR1 mRNA signal showed an early increase which levelled out in this line, whereas in 8-42 and UVM4 the increase was a late response (**Fig. 4B**). These observations based on RPKM levels agreed with the FC data in **Table 2** showing an early 10-fold induction of PSR1 in 8-27 PSR1 over UVM4 at d2/3 which decreased to 3-fold for d6. This compared with a constant 3-fold relative increase for 8-42 v. UVM4 over time (p-adj <0.05). In line 8-27, the transgene-specific YFP mRNA RPKM (**Fig. 4B**) matched the YFP antibody signal (**Fig. 4A**), indicating dependence of protein levels on mRNA levels.

### Expression patterns also placed target genes for PSR1 into four regulatory classes

The key gene expression changes (~50 genes out of OE-248) fell into four regulatory classes based on multivariate analyses of their expression patterns within our dataset. This outcome was in agreement with the above Venn comparison of the OE-248 list with the P-STRESS dataset, producing a similar result by independent means.

Target genes showing substantial expression changes with PSR1-OE are listed in **Tables 2** and **3** (full OE-248 list in data S1). Exemplar genes (summarized below) were taken from these lists including one or two of the best induced or repressed responses for regulatory classes I-IV (see above) and their gene expression levels (RPKM) are shown in **Fig. 4C, D**, alongside the data for PSR1 (**Fig. 4B**). Although these genes were all significantly linked to PSR1-OE (p-adj<0.05), their expression patterns were radically different from each other. This was investigated systematically by multivariate analysis (**Fig. 5A, B**) using the same OE-248 dataset as shown in the Venn analysis in **Fig. 3B** except that the full time course FC data was used instead of the magnitudes. This data-separation analysis identified the key genes in terms of (i) their differential expression strength (i.e. distance from origin); (ii) induction or repression (i.e. PC1, *x*-axis where approximately, induction *x*>0; repression x<0) and (iii) transgenic line (i.e. PC2, *y*-axis where data bi-plots cluster according to the algal line) (**Fig. 5A, B**).

**Fig. 5.**
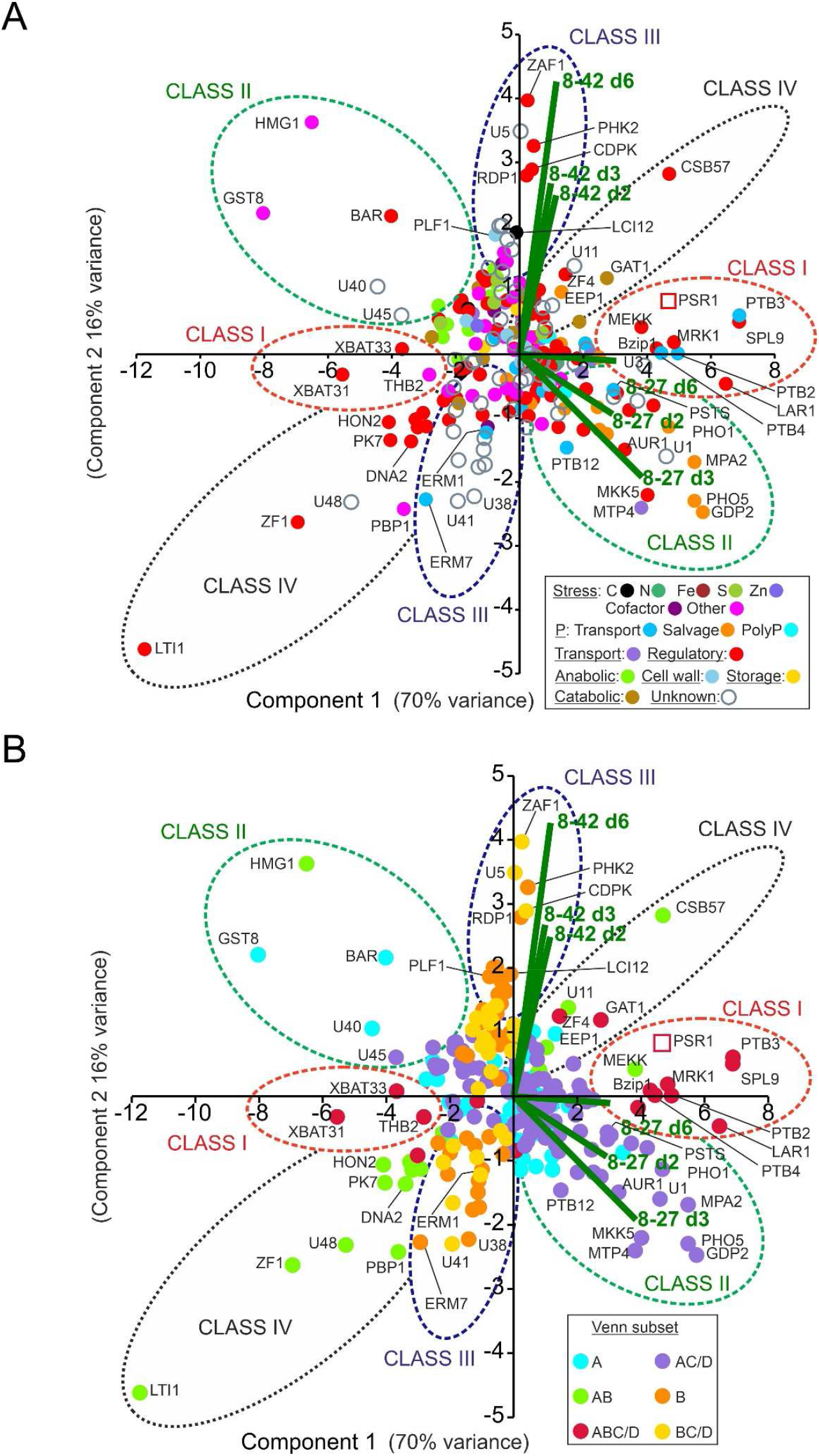
Multivariate analysis of PSR1-OE gene expression data. Shown are patterns of gene expression for OE-248 gene set (significantly regulated genes in line 8-27 or 8-42 relative to UVM4 with >2-fold cutoff). (**A**) Data points are color-coded for the 17 gene functional processes shown (inset). (**B**) The same chart is coded instead for the six Venn diagram sector subsets (inset) as described in **Fig. 3B**. In (**A**, **B**) the mean (n=3) relative gene expression time course data for the two transgenics v. UVM4 (log2(FC: fold-change) for d2, 3 and 6; i.e. 6 data points per gene) were analyzed by PCA. Biplots for each of these six data points are shown (-) and key gene expression changes are labelled. Clusters of genes are encircled and labelled Class I-IV. The data point for the complete PSR1 gene mRNA (not specific to endogenous gene or transgene) is shown (□).

The data were color-coded according to gene function processes (**Fig. 5A**) or the six Venn diagram subsets (**Fig. 5B**). In **Fig. 5A**, the genes displayed clustering according to gene function particularly differentiating P-transport (mid-blue) from P-salvage genes (orange) (**Fig. 5A**). In **Fig. 5B**, the genes were seen to fall into clusters which supported the four gene Classes I-IV derived from the Venn comparison with the P-STRESS dataset as shown in **Fig. 3B**. There were a handful of exceptions (e.g. GAT1, MEKK and HMG1) that could now be more accurately reassigned to different classes according to the PCA analysis (**Fig. 5B**). The findings for the four classes are summarized as follows:

**Class I** genes (Venn ABC/D) clustered with, or directly opposed, PSR1 gene activity, indicating a strong positive or negative correlation (FC data in **Fig. 5A, B**). This was supported by the high Pearson’s correlation coefficients for Class I gene to PSR1 (RPKM) (**Tables 2, 3**). This group also associated with PC1, which explained 70% of the variation, consistent with a strong influence. MEKK, a novel early-induced regulatory gene (**Table 3**), was re-assigned to Class I from the Venn AB set based on the PCA (**Fig. 5**). The P-stress induced transporter PTB2 (**Table 2**) was a highly expressed Class I gene (RPKM) (**Fig. 4C**). PTB2 was induced in proportion to the PSR1 mRNA levels (**Fig. 4C** cf. **Fig. 4B**) as implied by its association with PSR1 in the PCA (**Fig. 5A, B**). Also in Class I, transporter gene PTA1 (**Table 2**) showed a similar dependence of induction on PSR1 levels, although a temporal pattern of early induction followed by repression was seen in all lines (**Fig. 4C**). XBAT31 (**Table 3**) was decreased in proportion to PSR1 levels, where the temporal pattern was late repression (**Fig. 4D**).

**Class II** genes consisted of mostly P-scavenging genes that were highly induced in the strongest line only (8-27) (set AC/D) (**Fig. 5A, B**). The most dynamic “up” gene in the OE-248 dataset: PHO5/PHOX (an exophosphatase) (**Table 2**) showed transient induction in line 8-27, peaking at low PE levels but only to moderate RPKM levels compared with PTB2 (**Fig. 4C**). Regulated in a similar but less dynamic manner, was CAX1/VCX1 (putative PolyP synthesis gene) (**Fig. 4C**). There were 3 genes strongly in opposition to the 8-27 bi-plots (HMG1, GST8, BAR) and therefore, highly repressed in this line only (from sets A and AB) (**Fig. 5A, B**, **Table 3**). Irregular gene HMG1 was significantly increased in 8-42 but significantly decreased in line 8-27 (**Table 3**, **Fig 4C**). Originally placed in Venn subset AB, it was reassigned to Class II based on the PCA (**Fig. 5**).

**Class III** genes comprised highly induced/repressed genes associated only with line 8-42 (sets B or BC/D) including ZAF1, the most dynamic “up” gene in line 8-42, albeit late-expressed (**Fig. 4D, 5A, Table 3**). ZAF1 belonged to set BC/D and was therefore validated by its appearance in the P-STRESS dataset (although it was repressed here, **Data S1**), yet was unaltered in the stronger PSR1-OE line 8-27 (**Fig. 5**). ERM7 a Ca-dependent P-transporter channel was significantly repressed in line 8-42 only (**Table 2**) (**Fig. 5**). Class III genes included a high proportion of genes that were ectopically upregulated with P-stress in the absence of PSR1 in the P-STRESS data set (38% cf. 15% for the full OE-248 set) (Fig. S11, **Table 2 and 3, Data S1**).

**Class IV** genes consisted primarily of those from Venn diagram set AB, which showed equal responses in both lines (e.g. LTI1, ZF1, PBP1 and CSB57) (**Table 3**). In other words, they were equidistant from the two bi-plot clusters in **Fig. 5A-B** and showed similar expression data (FC) in **Table 3**. This was evident despite the differences in PSR1-OE levels. Although the GAT1 peptidase (**Fig. 4D**) and ZF4 were in Venn set ABC/D, they gravitated in the PCA towards Class IV (**Fig. 5B**). Functionally, GAT1 a peptidase, was not obviously linked to P-homeostasis (**Fig. 5A**).

### Genes that respond to stresses other than P-related are regulated differently

Further multivariate analyses revealed that genes for P-stress mitigation (e.g. P-transport, P-sparing or P-salvage) consisted of mostly early timing responses towards PSR1-OE whereas those associated with other stresses were late responses (**Fig. 6**). This allowed a hypothetical model to be drawn up including a feed forward mechanism for P-uptake (**Fig. 7**).

**Fig. 6.**
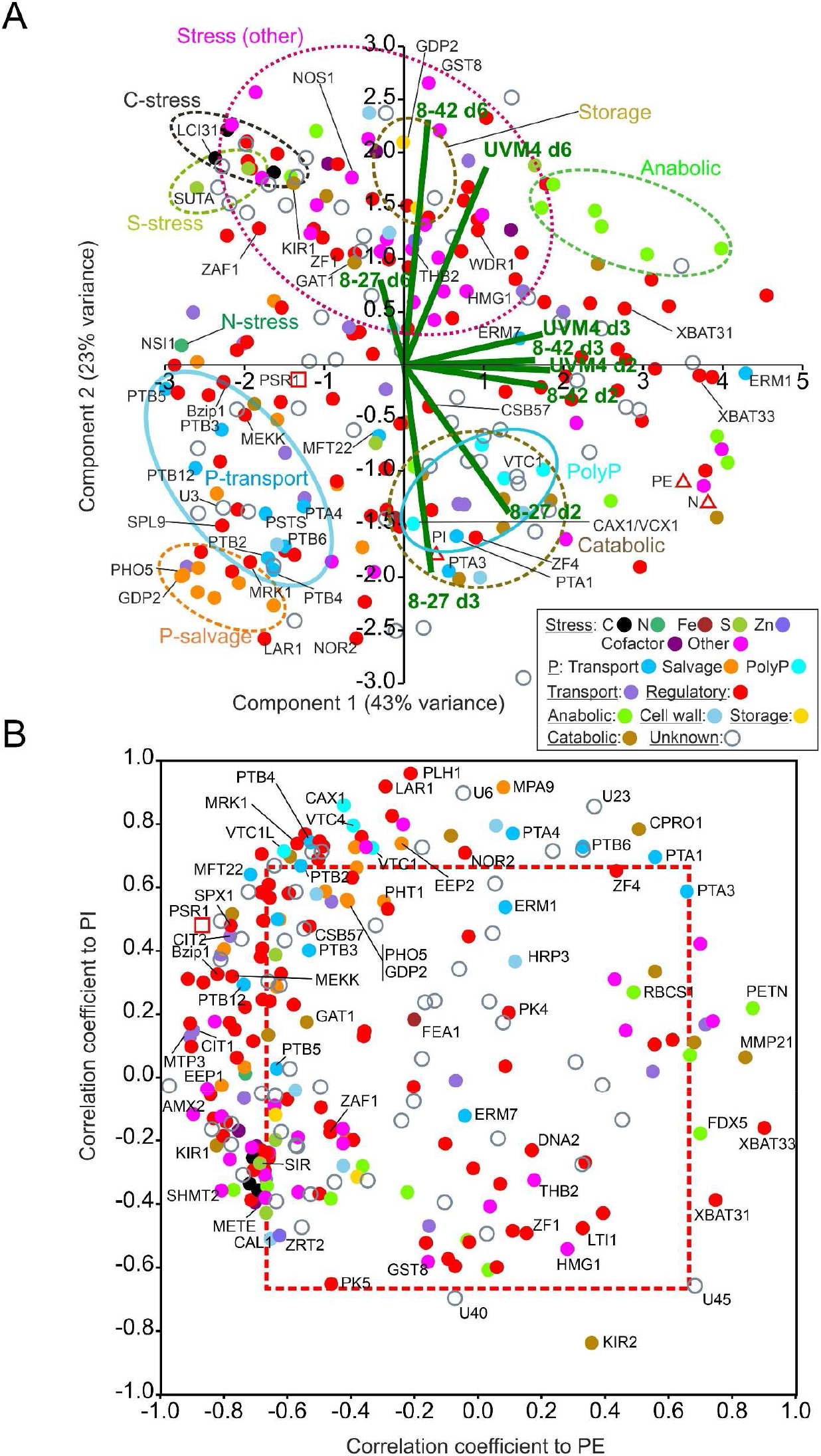
Temporal regulation of PSR1-OE gene expression. Patterns of gene expression focusing on temporal factors are shown for the OE-248 set. Data points are color-coded for the 17 gene functional processes shown (inset). (**A**) PCA analysis is shown for normalized mean (n=3) RPKM data along with the PE, PI and N measurements (Δ). The biplots for the nine data sets (UVM4 control and both lines: time points d2, 3, 6; i.e. 9 data points per gene) are shown (-). Clusters highlighting one specific gene functional process are encircled. (**B**) Plots of Pearson’s correlation coefficients for RPKM data v. PI and v. PE for each OE-248 gene. Coefficients outside the boxed region (---) were significant for PI or PE (P<0.05). In all cases mean data was from n=3 culture replicates.

**Fig. 7.**
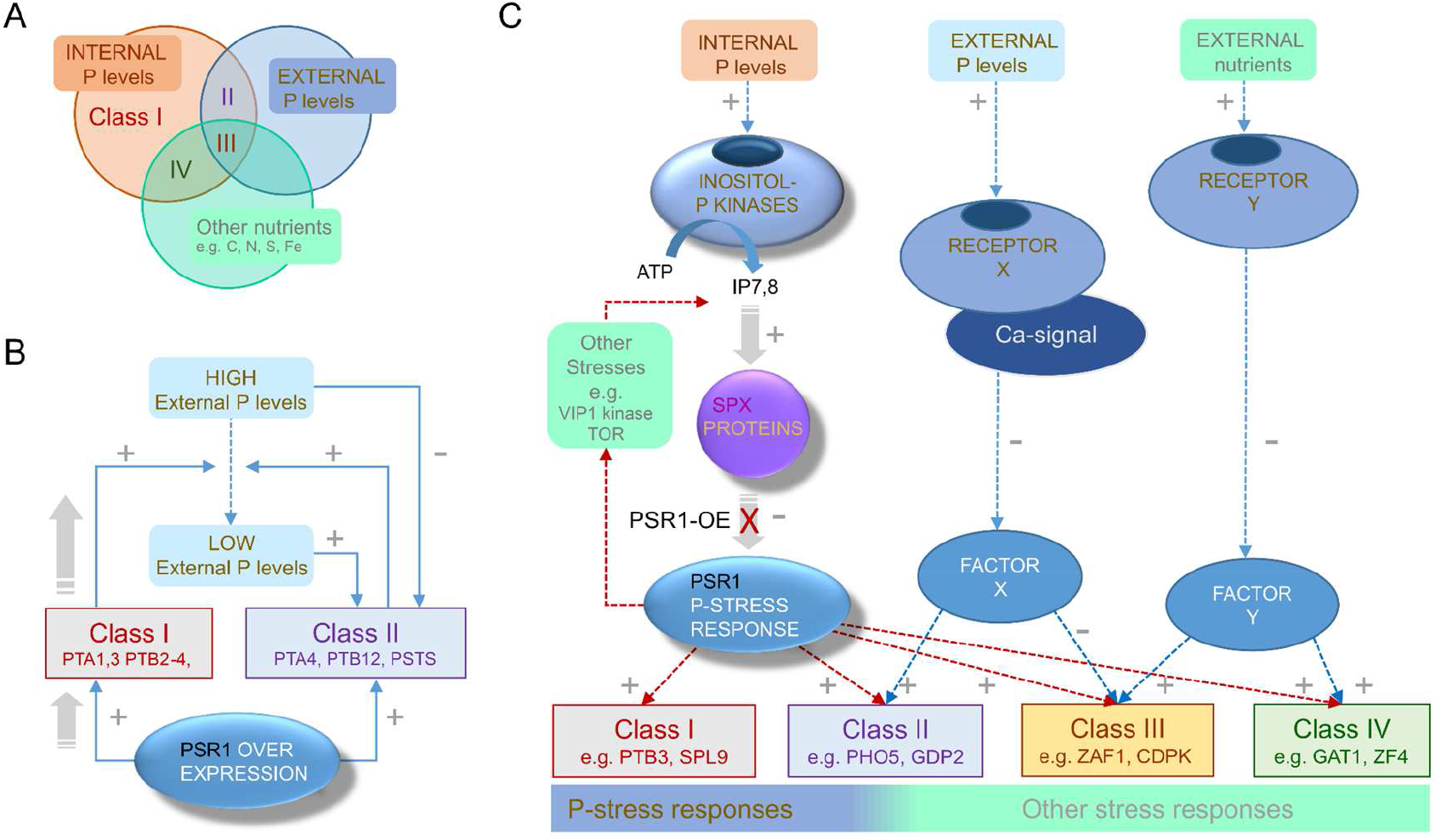
Proposed regulatory networks for PSR1-dependent genes. (**A**) Venn diagram illustrating the three nutrient factors or stresses that appear to influence the four Class I-IV subdivisions of the OE-248 set of genes that show significant expression changes (classes defined in **Fig. 5A, B**). Each Class I-IV is shown occupying a sector in the Venn diagram indicating the principal nutrient factor or stress that is proposed to influence it. (**B**) Proposed feed-forward mechanism explaining enhanced P-uptake driven by PSR1-OE on Class I genes under initially high PE conditions. Activation of Class I P-transporter genes (+) reduces the PE levels which are inhibitory (-) towards Class II genes. Subsequent activation of Class II genes which include P-transporters, further reduces PE. (**C**) Proposed signal network for transducing changes in PI, PE and other nutrient factors or stresses (e.g. C, N, S, Zn etc.) towards gene induction for the four different regulatory Classes I-IV. Class I genes are regulated principally by PSR1-mediated perception of PI and can be activated by PSR1-OE in the presence of high PE. Class II genes are PSR1-dependent but require agreement between PI and PE sensing mechanisms. Class III are PSR1-dependent but are inhibited by low PE and activated by low levels of other nutrients or stresses. Class IV are PSR1-dependent but activated by low levels of other nutrients or other stresses and are unaffected by PE levels.

To probe the entire OE-248 dataset, expression change magnitude was factored out to focus on temporal factors using normalized RPKM data (Data S1). This also allowed co-analysis with physiological measurements (PI, (internal Pi), PE (external Pi) and N (medium N)). Both PCA (**Fig. 6A**) and correlation coefficient analyses (**Fig. 6B**) clustered genes into discrete functional categories. For instance for the PCA: S-assimilation (2), C-assimilation (3), storage (2), putative PolyP synthesis (4) and P-salvage (8) functions and other stress-related functions (18 genes). In **Fig. 6A**,virtually all the genes for P-stress (early induction) were separated from those for all the other stress responses (N, S, C, Fe, Zn, Cofactor, motility, DNA-repair etc.) which were late induced (d6) (**Fig. 6A**, fig S12A). The latter group amounted to mostly weaker changes (signaled by their closer clustering to the origin in the earlier PCA in **Fig. 5A**) but with some exceptions (e.g. HMG1 and GST8), which underwent larger changes (**Fig. 5A** cf. **Fig. 6A**). Along the PC1 axis, PSR1, P-transport and P-salvage genes (left) were separated from those PolyP synthesis, which associated closely with internal P levels (PI) (bottom right, **Fig. 6A**). Repressed genes in opposition to PSR1 (e.g. XBAT31, 3) associated with PE or N (on the right, **Fig. 6A**).

In **Fig. 6B**, correlation coefficients for P-levels: PE and PI, were plotted for the gene RPKM data. A strong bias was seen towards genes that clustered with PSR1. PSR1 itself showed a close negative PE correlation (p<0.05, **Table 2**) and a weak positive PI correlation (NS, **Data S1**). These genes were primarily linked to line 8-27 (Venn subsets ABC/D, AC/D or A) i.e. Class I and II (fig S12B). A few genes showed regulation in the opposite direction (e.g. FDX5, XBAT31, 3; also Class I and II) which were repressed by PSR1 and correlated positively with PE (**Fig. 6B**, fig S12B). A significant positive correlation was seen with PI for the four putative PolyP synthesis genes along with PHL1, a nucleotide triphosphate hydrolase and regulatory genes such as LAR1 (**Fig. 6B**) (**Data S1**). Many (13) Class IV genes (Venn AB) correlated with PI only (e.g. NOR2 P<0.05, rest NS). Some (5) correlated with PSR1 and/or PE (**Fig. 6B**, fig S12B; note MEKK re-assigned to Class I).

## Discussion

The ability to switch on luxury-P uptake would greatly facilitate microalgae-based wastewater treatment. Achieving hyperaccumulation of P even when the element is plentiful would avoid the need for stress preconditioning. This would simplify the process of fitting algae into the existing wastewater plants (*3*). An important question was whether this could be achieved by increasing the levels of a single regulatory gene. To test this, we over-expressed PSR1, a global regulator of the P-stress response, generating a spectrum of PSR1 over-expression levels. In the strongest over-expresser line (8-27), P-uptake rates were increased 3-fold and maximum biomass P levels, stored as PolyP, were increased 4-fold to 8%DW. Complete removal of culture P was brought forward by at least 3 d during batch culture where the control strain had removed only about 70% after 7d. This was achieved by late log phase at relatively low biomass concentration (0.2 mg DW L^-1^), uncoupling P-uptake from biomass production.

With biomass P at 8%DW and no P left in the medium however, further growth was accompanied by a near complete remobilization of PolyP reserves back to control levels. This remobilization of P was probably inevitable since it would be required to sustain 2-3 further cell divisions to reach stationary phase at the same rate as the control (*10*). A better understanding of the genetic mechanisms of PSR1 action could help to address the transience of PolyP reserves, a potential shortcoming.

We found that during batch culture of the control strain (UVM4) in replete medium (TAP), only minor reductions in nutrients were associated with significant gene expression changes but Phosphate Stress Response genes (*16*) were under-represented compared with other nutrient stress genes (C, S, N, Fe etc.). Early reductions seen in protein translation gene expression may have represented a general response, including P-stress. C-stress was a prominent factor in our UVM4 data and has recently been shown to lead to a TOR-kinase linked drop in key amino acids in *C. reinhardtii* (*21*), probably impacting translation.

Against this backdrop, P-uptake levels were drastically altered to different degrees in different transgenic PSR1 over-expresser lines. Evidence strongly pointed towards over-expression of PSR1 as driving the increased P-uptake levels. For instance, PSR1 gene mRNA levels were found to have a strong negative correlation with external P levels (PE) and were in proportion to P-uptake rates in PSR1-OE lines. Transgenic over-expression of PSR1 increased the endogenous wild type PSR1 gene mRNA levels and also mimicked the induction of a cohort of genes associated with reported P-stress (*13, 16*) and P-resupply (*22*). Altogether 248 genes were found to be significantly altered in gene expression relative to the control (UVM4). Of these, 60% (146) were also seen in one univariate P-stress experiment (*16*) (but not necessarily described) and the remaining 40% (102) were novel. Despite these similarities, there were major differences in the way four different sub-classes of these genes were affected by PSR1 over-expression. These differences were associated with specific gene functional processes implying that different environmental cues (internal and external P levels and other nutrient stresses) were modifying the gene responses to varying extents (**Fig. 7A**).

### Proposed feed-forward model accounting for the rapid P-uptake

Evidence suggested that a relatively small group of genes (Class I) were driving the expression of a larger set of genes (Class II) by a feed-forward loop mechanism (**Fig. 7B**). This was inferred from multivariate analyses and the timing of specific genes. For instance Class I genes, consisting of P-transporters (PTA1, 3, 4; PTB2-4) and several putative regulatory genes (e.g. LAR1, SPL9, Bzip1, MRK1, MEKK) were elevated early, in the presence of high external Pi levels (PE). These genes were upregulated in more than one transgenic PSR1 over-expression line and were strongly associated with PSR1 expression levels either by correlation (**Tables 2, 3**) or PCA (**Fig. 5A, B**). PSR1 RNA levels in turn were proportionate to P uptake rates in the 3 lines (UVM4, 8-42 and 8-27) and also inversely correlated well with PE.

The timing of putative P-transporter genes supported the model, whereby low P-affinity transporter genes were progressively induced and supplanted by those with higher affinity or active transport by PSR1-OE: 1^st^ PTA1,3,4 (d2 peak, low affinity; Class I), 2^nd^ PTB2-4 (d2-d3 peak, high affinity; Class I) and 3^rd^ PSTS (active transport: a putative subunit of a prokaryotic type ATP BINDING CASSETTE (ABC) transporter) and PTB12 (high affinity) (both d3 peak; Class II) (**Table 2, Data S1**) (*17*). Of the PTB genes, only Class I PTB2-4 (cf. PTB5, 6) showed abundant transcript levels (RPKM) as well as high FC’s and were presumably the key players (**Fig. 4, Data S1**). The induction of enzyme activity levels for PTB (higher P-affinity) has been noted with P-stress (*17, 23*)

The model proposes that most of the P-stress related genes (Class II) were inhibited by PE, until levels had been sufficiently reduced by the action of the small group of Class I genes (**Fig. 7B**). This is supported by the behavior of the PHO genes which encode P-scavenging exophosphatases repressed by P (*18*). In our multivariate analyses (**Fig. 5A, B**) these co-clustered with a large group of PSR1-dependent genes responsible for much of the P-stress response characterized as Class II. These genes tended to peak in expression after the transient apogee of P-biomass, for instance one of the strongest relative gene expression (FC) changes was PHO5/X (Class II) which peaked at d3 whereas PTB3,4 peak at d2 (Class I) (**Table 2**). PHO5/X is also one of the most strongly altered genes in P-stress experiments (*16, 18*). The feed-forward aspect of the model is also supported by the induction of Class II putative P-transporters (PSTS, PTB12) which could reinforce reductions in PE driven by Class I genes. This could provide an explanation for the rapid P-uptake seen in the transgenic lines, along with complete removal of P.

Assembling a gene model of the P-stress sensing apparatus integrated the findings in this work with knowledge in the literature, revealing gene targets for improving replication of the luxury P-uptake response (**Fig. 7C**). The model incorporates the well-known Inositol PolyP (InsP7, 8) pathway, which is presumed to operate in *C. reinhardtii* for the detection of internal Pi (*24*). It is proposed that for Class I genes, increasing PSR1 levels overcomes InsP7, 8-mediated inhibition that is signaling an ample P-supply, perhaps by titrating out the SPX1 protein. In vascular plants the PSR1 homologue PHR1 co-ordinates multiple aspects of the response to low phosphate (*25*). Under high P it is prevented from binding its target site in the promoter of downstream genes by inositol P dependent binding to SPX1 (*26, 27*). In the current study the SPX1 gene was itself increased by PSR1 over-expression but this was not an early response and was also relatively weak (**Table 2**). In this model, PSR1-dependent Class II genes (e.g. PHO5/X) are regulated by this pathway as well as a PSR1-independent pathway for sensing P-levels (**Fig. 7C**). This would render the genes initially insensitive to PSR1 increases until PE levels decreased. Intriguingly, an external P-receptor has been proposed in diatoms on the basis of Ca-mediated signaling for P (*28*) culminating in gene expression changes though unknown means (Factor X) (**Fig. 7C**). This mechanism could also explain the occurrence of a minority of PSR1-independent genes that are reportedly altered in P-stress (Fig. S11, Data S1) (*16*). According to the model, these would be regulated by the external P sensor pathway only. Additionally, the TOR signaling complex, which integrates nutrient status to control growth and various anabolic or catabolic pathways, has been placed downstream of PSR1 (via LTS8) (*29*). TOR also interacts with the VIP1 kinase which produces InsP7,8 so could potentially act upstream of PSR1 via SPX1 (**Fig. 7C**) (*30*).

### Mg^2+^ as a dynamic counterion for PolyP accumulation

Achieving a greater retention of the high P in biomass levels could be desirable, even if this incurred a growth penalty; providing an incentive to explore the underlying molecular changes involved in uptake and remobilization of PolyP. High P-uptake driven by ectopic PSR1 increases was associated with a transient accumulation of very large PolyP granules. This was closely associated with a rapid uptake of Mg^2+^, suggesting that this cation, rather than Ca^2+^, was acting as a principal counter-ion in PolyP storage under these circumstances. Both are the predominant cations associated with PolyP in *C. reinhardtii* and higher measured levels of Mg^2+^ cf. Ca^2+^ have been reported in isolated PolyP granules (*31*) but in another report, they were roughly equal (*32*). We noted a 2:1 molar ratio of P: Mg^2+^ for the uptake rates (R_MAX_) (**Table 1**). This lent credence to the idea that Mg^2+^ was the sole counterion, since one divalent cation would bind two P groups in the polymer, given that each P group carries one negative charge (OH^−^) (*6*). Although a shortage of Mg^2+^ might explain the instability or diffuse appearance of enlarged PolyP granules, Mg^2+^ levels were not actually depleted beyond a low baseline of 3 mg/L, whereas P was depleted below detection.

### Accounting for the transience of stored PolyP

Another factor that could have impacted PolyP stability was the transcriptional control of the synthesis pathway. Several Class II genes were putatively associated with PolyP production, for instance the CDF family cation transporter MTP4 (35-fold) and Chromate ion transporters CIT1, 2 (5-fold) which could be responsible for the rapid Mg^2+^ uptake. However, a Ca^2+^/H^+^ vacuolar antiporter (CAX1/VTC1) along with three putative PolyP synthesis genes (VTC1, VTC1L and VTC4) were also induced but by lower levels (2-3 fold). Interestingly, a triphosphate hydrolase (PLH1) (6-fold) co-clustered with this latter group of genes, where all exhibited a close positive correlation with internal Pi levels).

It was important to note that many of the Class II genes showed transient expression peaks with PSR1 over expression, and this could contribute to PolyP remobilization. The sharp decrease (particularly evident for genes such as PHO5/X) could be due to a negative feedback mechanism and/or a high sensitivity to decreases in PSR1-YFP protein levels. The latter possibility was supported by use of protein translation inhibitors in relation to PHO5/X (*16*). Once P was removed from the medium, a drop in PSR1-YFP fusion protein was evident in both transgenic PSR1-OE lines, particularly striking in the case of the strongest OE line. This points towards eventual post-translational down-regulation of PSR1-YFP protein levels, as a contributory factor, towards the end of the experiment because late decreases in the transgene mRNA were only moderate (down 50%) and only seen in the strongest line. In the transgene construct, expression was driven by a *C. reinhardtii* PSAD light-regulated promoter. The wild-type PSAD gene levels dropped by 35-40% over the experiment (**Data S1**) but this was seen in all lines so it cannot account for changes that occurred in one line only. The endogenous PSR1 gene continued to increase in expression, as external P was depleted, in all lines (**Fig 4B**). In the literature however, upregulation of the PSR1 gene mRNA has been shown to be transient in univariate stress experiments (P, N, S) (*14, 16, 19*).

There was further evidence from our work and others that might also account for the transience of some Class II gene expression. For instance, THB2 (a Class I regulatory gene) was late-repressed by PSR1 over expression. This truncated hemoglobin is thought to act as an NO scavenger (*33*). In addition, there was also late induction of NOS1 (3-fold) a possible flavodoxin/nitric oxide synthase with a ferrodoxin reductase-type FAD-binding domain. Together these changes were consistent with observed increases in NO associated with P-stress (*33*). Switching off THB2 reportedly represses P-stress induction of genes such as PHO5/X thus providing a possible negative feedback mechanism accounting for the transient nature of many of the Class II genes as well as PolyP (*33, 34*). Poly P synthesis and mobilization are also regulated directly at the enzymatic level by inositol phosphates (InsP7, 8) binding to SPX domains attached to VTC proteins and PTC1 (**Fig. 7C**) (at least in yeast) (*35, 36*). Therefore, PolyP remobilization and lack of PolyP synthesis could be transduced by this mechanism in the absence of PE (*35, 36*).

### PSR1 influences ‘*other nutrient stress*’ genes and general stress responses

The global nature of PSR1 gene regulation beyond P-homeostasis has been indicated (*19*). Likewise, new roles for PolyP production in mitigating other stresses (e.g. S) through utilization of excess ATP have been noted (*37*). In this context, we found that PSR1 over-expression also led to changes in genes that could alleviate other nutrient stresses (C, N, S, Zn, Fe and vitamin cofactors) or heralded general stress responses in *C. reinhardtii* (e.g. DNA-repair, reproduction, motility, defense, storage products etc.). Most of these genes were regulated later than the P-stress genes however, and generally the responses were weaker but with exceptions (e.g. GST8, HMG1, LTI1 and GAT1). Some of these genes fell into **Class II** (possibly affected by PE), for instance GST8 a glutathione S-reductase (oxidative stress) and HMG-CoA reductase (HMG1) that were both strongly repressed with PSR1-over-expression. Reduction of sterol synthesis (i.e. ergosterol) could be a consequence of repressing the reductase, and this is a known stress response in other organisms (*38*). The GAT1 peptidase (early response) may act to alleviate N-stress and a potential drop in key amino acids as reported for C-stress (TOR-kinase mediated) (*21*).

Many of the *‘other nutrient stress ’* genes fell into **Class III or IV** type regulation where nutrient factors other than P were proposed to impact gene expression (**Fig. 7C**). For instance, in Class III (e.g. S-stress: SUTA and SIR; low-C stress: LCI12), genes were paradoxically strongly regulated with weaker PSR1 over-expression increases and these were generally late responses. Induced genes responsible for cofactor auxotrophy such as METE (bypasses a requirement for vitamin B12) also fell into this class (*39*). Interestingly, the aforementioned S-stress function genes induced by PSR1 over-expression were not among the most strongly induced genes reported for a univariate S-stress experiment (*40*), whereas ATS1, which was induced strongly in this report showed significant repression in our work (**Table 3**). Therefore, a somewhat different complement of genes was being influenced by PSR1 over-expression in our work.

A very high proportion of genes in **Class III** (38%) showed reported enhanced expression changes with P-stress in the absence of PSR1 (ectopic change) (Fig. S11, Data S1) (*16*). This was mostly ectopic induction where presumably PSR1 opposes induction and here, the proposed PSR1-independent P-stress mechanism might be responsible for the increase. We note that conversely, the Class III ZAF1 gene was repressed in a PSR1-independent fashion by P-stress (Fig. S11, Data S1) (*16*) but we found it strongly induced in the weak PSR1-over expression line. Therefore, it could instead be induced by PSR1 but repressed by a PSR1-independent P-stress mechanism (**Table 3**). We propose that in Class III there might be a conflict between the two (or more) mechanisms of P-stress perception that could act to shift the focus more exclusively to P-homeostasis genes when this particular stress dominates (**Fig. 7C**). This is consistent with our observed dampening down of Class III gene expression changes when PSR1 is over-expressed at higher levels, perhaps because PE is reduced faster.

Much of the **Class IV** gene expression did not correlate with PSR1 levels or external P and might depend more on other nutrient stresses (represented in **Fig. 7C** by unknown Factor Y). With Class III and IV, late responses prevailed therefore PSR1-dependent control might be exerted via an intermediary gene, perhaps one of the regulatory genes in Class I, and this would permit complex modes of regulation.

## Conclusions

In this study we demonstrated that over-expressing a single gene (PSR1) led to a remodeling of metabolism that induced luxury P uptake. We found that enhanced P-uptake was accompanied by an accumulation of large PolyP storage granules. Our data strongly implicated Mg^2+^ as the principal counter ion for P-storage in this form. We also identified a possible feed-forward mechanism where ectopic PSR1 promotes induction of a small set of *‘driver genes’* at high P-levels. These go on to reduce external P levels, allowing the induction of P-repressed genes responsible for further P-uptake so that luxury uptake of P was induced. As expected for a transcription factor, RNAseq analysis showed key genes were induced/repressed. Clearly there are other regulatory mechanisms that might operate at post-transcriptional levels but our results provide a foundation to build on our proposed models. Although only one out of three transgenic lines showed strong PSR1 over-expression, this was a strength that allowed correlation of responses to levels of PSR1 expression. It could also be seen as a limitation, however published corroborative data was used to compare and substantiate findings. We carried out over-expression of PSR1 in only one engineered *C. reinhardtii* UVM4 strain lacking a cell wall and aspects of gene silencing to favor transgenic expression (*41*). Like many *C. reinhardtii* strains it carries *Nit1, 2* mutations preventing use of nitrate (*20, 42*). Nevertheless, recent data suggests that these potential problems can be overcome to give high productivity in UVM4 (*43*). Furthermore, our technology is also likely to be transferrable since a related *Chlamydomonas* sp. has shown good growth on wastewater (*44*) and PSR1 homologs carrying out P-starvation responses have been found in diverse taxa, such as the marine diatoms and higher plants (*27, 45*). Although we found hyper-accumulation to be transient, semi-/continuous cultures could be employed to hold the culture at peak biomass P. Alternatively, the moderate over-expresser lines also showed slower remobilization. Turnover could also be blocked by stacking mutations in PolyP remobilization genes such as *PTC1* (*35, 36*). Collectively, these results will accelerate progress towards circular bioeconomy algal solutions for wastewater treatment or eutrophic waterbody bioremediation.

## Materials and Methods

### Algal strains, culture and harvesting

*C. reinhardtii* strain UVM4 (*20*) and its derived transformants were cultured in an Algaetron AG230 (Photo Systems Instruments, Czech Republic). The cells were grown in Tris Acetate Phosphate (TAP) media without Na_2_SeO_3_ (*46*) in 100μmol photons m^−2^ s^−1^ constant light at 25°C. For the microscopy, cell lines were grown in bijou containers for 3-4 d without shaking. For the growth experiments in batch culture, liquid cultures were inoculated from a 3-4 d starter culture for an initial OD_750nm_ of 0.005 in 250 mL conical flasks, which were shaken at 150 rpm and growth was monitored for 7 d. The transgenic lines *C. reinhardtii* LC8-27, LC8-42 and LC8-2 were grown in parallel with a non-transformant control strain (*C. reinhardtii* UVM4), in triplicate. Samples (1-30 mL) were centrifuged at 5000 g for 10 min and the supernatants were filtered (22μm pore size Millex-GP Syringe Filter Unit, Merck) and frozen at −20°C for measuring medium composition. The biomass pellets were washed twice with deionized H2O, flash frozen with liquid nitrogen and stored at −80°C.

### Generating the PSR1-OE construct and the transgenic lines

The transgenic lines *C. reinhardtii* LC8-27, LC8-42 and LC8-2 were independent transformants designed to over-express the PSR1 gene. Constitutive over-expression of the PSR1 gene (PSR1-OE) in algae was driven by the PSAD gene promoter of *C. reinhardtii*,followed by the RBCS intron, to maximize expression. The DNA construct was assembled as follows: the PSR1 gene (Cre12.g495100.t1.1) with an inserted 3xHA-tag was synthesized (Genscript Biotech Corporation, UK) and cloned into pUC57 via the *StuI* restriction site. This plasmid was then used as a template for Golden Gate-based cloning (MoClo Plant Kit, Addgene) (*47*). The following level 0 plasmids were used: pCM0-001 (PSAD prom), pCM0-024 (RBCS2 intron), pCM0-044 (mVenus, incl. Strep-tag), pCM0-114 (PSAD term) all from (*47*) and L0_PSR1 (PSR1 cloned into pAGM1287 (MoClo Plant Kit) in this study). For the generation of level 2 plasmids for *C. reinhardtii* transformation, the following level 1 plasmids were used: pAGM4673 (L2 backbone, MoClo Plant Kit), pICH41822 (L2 end-linkers MoClo Plant Kit), pICH54011, pICH54022, pICH54033, pICH54044 (Dummies, MoClo Plant Kit), pCM1-27 (ParoR,) (*47*) and L1_PSR1 (PSADprom-RBCS2intr-PSR1-mVenus-PSADterm, this study). All restriction/ligation reactions were performed using BpiI (BbsI) (Fisher Scientific) or BsaI-HF (NEB) together with T4 ligase (NEB) in a total volume of 20 μL containing 1x BSA, 1x T4 ligase buffer, 5U restriction enzyme, 200U T4 ligase. The typical ratio between destination plasmid and entry plasmid/parts was 1:2, using 75 ng of the acceptor plasmid. Level 1 assembly reaction: 20 sec 37°C, 26x (3 min 37°C, 4 min 16°C), 5 min 50°C, 5 min 80°C, hold 16°C. Level 2 assembly reaction: 45x (2 min 37°C, 5 min 16°C), 5 min 50°C, 10 min 80°C, hold 16°C. All plasmid concentrations and quality were determined using a NanoDrop (ND-1000, Labtech). Correct assembly was confirmed by sequencing (Genewiz). Primers and DNA constructs are listed in Tables S1, 2.

### Genetic transformation of *C. reinhardtii*

For *C. reinhardtii* transformation, UVM4 was grown in liquid TAP for 2 days until mid-logarithmic phase (1-4 x 10^6^ cells/mL). Cells were collected by centrifugation (2500 g, 10 min) and the pellet was washed twice with ice-cold EP buffer (electroporation buffer: 40 mM sucrose, 10 mM mannitol and 10 mM CHES pH 9.25). The pellet was resuspended in EP buffer to a final volume of 1 x 10^8^ cells/mL. Transformation was performed by electroporation using a NEPA21 (Nepa Gene Co. ltd.) and 0.2 cm cuvettes (Nepa Gene Co. ltd.) at the following settings: 2x poring pulse 300 V, 4 ms length, 50 ms interval, 10% decay rate, polarity +; 1x transfer pulse 20 V, 50 ms length, 50 ms interval, 40% decay rate, polarity +/-. For each transformation, 25 μL cells (2.5 x 10^6^ cells) and 5 μL Plasmid-DNA (500 ng) were used. After transformation, the cells were kept at dim light (2-3 μE) for 16 h and plated on fresh TAP plates containing 10 μg/mL Paramomycin (Sigma). Plates were incubated for 10 d at 30-50 μE constant light. Colonies were picked and sub-cultured weekly, for 3 weeks in liquid TAP + Paromomycin. Surviving colonies were screened by Colony PCR where 100 μL of cell suspension of each colony was collected by centrifugation (15,000 g 1min). Pellets were resuspended in 50 μL 5% Chelex-100 (Sigma). Samples were boiled for 10 min, cooled down on ice, vortexed and centrifuged again. 1 μL supernatant in a total reaction volume of 20 μL was used as template and PCR was performed using Q5 polymerase (NEB). Primers are listed in **Table S1**. Stably transformed lines were screened by western blotting for levels of intact fusion protein.

### Protein extraction and Western blot analysis

Total protein was extracted according to (*48*) and the equivalent of 10 μg of chlorophyll was loaded per lane on a Tris-Glycine based SDS-gel (Mini-Protean®TGX, Bio-Rad or Novex™ 8%, Thermo Fisher) and transferred onto a PVDF membrane using the Trans-Blot®Turbo™ Transfer System (Bio-Rad). Chlorophyll was determined as below. The membrane was blocked for 1h in 5% (w/v) skimmed dry milk in TBS-T. To reduce background cross reaction the primary anti-GFP antibody (1:5000, ab6556, Abcam) was preincubated with a membrane containing protein extract of UVM4 before being added in 3% milk TBST for incubation overnight at 4°C. The membrane was washed 3x in TBST for 10 min and incubated with the HRP-conjugated secondary anti-Rabbit antibody (1:5000, 111-035-144, Jackson Immuno Research) in 3% milk TBST for 1 h at room temperature. ECL detection was performed using the SuperSignal™ West Pico Chemiluminescent Substrate (Thermo Scientific).

### Chlorophyll measurements

Chlorophyll concentration was determined by pelleting 0.1-1mL *C. reinhardtii* cell culture (max. speed, 10 min) and resuspending the pellet in 1mL 80% acetone in MeOH. After a second centrifugation step for 5 min, absorbance was read at 663.6 nm, 646.6 nm and 750 nm using a spectrophotometer (Jenway 6715UV/Vis, Geneflow) (*49*).

### Confocal Microscopy

Culture samples of 200 μL were collected and 2 μL of DAPI stain (1 mM) was added, and samples were incubated in the dark for 4 h. For nuclear targeting, live cell images were captured with a Zeiss LSM880 + Airyscan Inverted Microscope (Carl Zeiss) using a Plan-Apochromat 40x/1.4 Oil DIC M27 objective. Filters were set as follows: Venus Ex. 514 nm, Em. 520-550 nm; DAPI Ex. 405 nm, Em. 420-475 nm (DAPI-DNA) and 535-575 nm (DAPI-polyP). Chlorophyll autofluorescence was captured with the 514 nm laser at 670-720 nm. Visualization of PolyP from the time course experiment was carried out as above with the following differences. After DAPI incubation, samples were fixed with glutaraldehyde: 25% (SIGMA) stock was added at 20μL/mL of culture and incubated for 20 min before being flash frozen with liquid nitrogen and stored at −70°C for later analyses. A Plan-Apochromat 63x/1.4 Oil DIC M27 objective was used (Zeiss LSM880+ Airyscan Upright Microscope, Carl Zeiss). Time course DAPI-polyP images were obtained as 6-8 Z-stacks and further processed as a Z-projection using the software Fiji (Image J) (*50*).

### Analysis of medium composition

Filtered supernatant samples were diluted 5-50 fold in dH2O and tested for soluble phosphate concentration (PO_4_^3-^-P) according to (*51*). Ammonium concentration in filtered media samples (NH_4_^+^-N) was determined using the Hach^®^ cuvette test LCK-304. Samples were diluted 50x-100x, due to interference by components of TAP media. IC measurements were made as follows: anion and cation analyses were performed using an ion chromatographer (Metrohm 850 Professional IC), with an 896 Professional Detector. Sterile filtered supernatant samples were diluted between 10-20X. The anion pump injector used 20 μL of the diluted sample and was analyzed with a Metrosep A Supp 5-150/4.0 separation column (flow rate of 0.7 ml/min). The cation pump injector used 10 μL of the diluted sample, which was analyzed with a Metrosep C4-100/4.0 separation column (flow rate of 0.9 ml/min).

### Biomass composition analysis

For phosphate in biomass, pellets were dried under vacuum using a SpeedVac Plus (SC210A - Thermo Savant Instruments) overnight, and dry weight was determined. A second drying period (overnight) ensured that dry weight data was accurate. The dry pellets were digested with an oxidizing reagent at 100°C for 60 min, using a Hach Lange LT200 Dry Thermostat (*51*). The digested samples were diluted 10X-25X and tested as mentioned above.

### RNA extraction and sequencing

RNA was extracted by grinding frozen algal pellets in liquid nitrogen, followed by extraction with a Qiagen plant RNA mini kit (Qiagen). Subsequent RNA sequencing work was carried out by the Next Generation Sequencing Facility (Leeds Institute of Biomedical & Clinical Sci.). RNA quality was checked using a 2100 Bioanalyzer and Expert software (Agilent). 100 ng total RNA of each sample was used to generate a TruSeq stranded RNA Illumina compatible library from which rRNA was removed using rRNA-specific depletion reagents. After size selection and adaptor removal with AMPure beads (Beckan Coulter), library concentrations were determined by qPCR before combining to make an equimolar pool that was sequenced (75bp single end sequencing read HiSeq3000 lane; Agilent; Santa Clara, USA).

### RNAseq analysis

RNAseq data was processed using software on the Galaxy Server (https://usegalaxy.org) except as noted. The reference genome and gene annotation files were obtained from JGI (https://genome.jgi.doe.go). Sequence data were checked for lower quality bases and adaptor sequences with FastQC (https://www.bioinformatics.babraham.ac.uk/projects/fastqc/) before and after trimming using Trimmomatic. The trimmed sequencing reads were aligned using Hisat2 to the reference genome file Chlamydomonas_reinhardtii_JGI_v5.5. Reads were counted with FeatureCounts using the gene annotation file Creinhardtii_281_v5.5.gene_exons.gff3. For determining gene expression levels, these counts were converted to RPKM in Microsoft Excel, normalizing to gene length and total reads (Data S1). To obtain relative gene expression data values (Fold change: FC) as log2 (FC) with associated significance (P-adj values), FeatureCount files for replicate (n=3) cultures were compared for experimental v. controls using DESeq2 (Data S1). For determining RPKM for elements of the transgenic construct (e.g. YFP) or wild-type specific components of the PSR1 gene (e.g. 3’UTR) in the genome, trimmed files (Trimmomatic) were converted to FASTA files (Fastq to Fasta converter). The sequences within were renamed numerically with Rename sequences (numeric counter) and a blast dbase created for each file using Makeblastdb(nucleotide) before carrying out blastn with the appropriate gene fragments to obtain the counts, which were converted to RPKM as above.

### Further transcriptomic data analyses

Version 5.6 gene annotation data for *C. reinhardtii* was downloaded from JGI (https://genome.jgi.doe.go) and assigned to the curated transcriptomic data in Microsoft Excel. Time course data was processed as follows: for investigating changes in the control UVM4, relative gene expression data was derived for d6 v d2 (late) and d3 v d2 (early) as above. In each case genes were ranked in Microsoft Excel by the up and down values to obtain four sets of significantly regulated genes (>2-fold P-adj<0.05). Panther gene ontology codes from the annotation were used to analyze gene function for the four gene sets at http://www.pantherdb.org/. In addition, the top 200 ranked genes by FC for each set were manually curated into gene functional roles (and unknowns) which were formulated to match the specific requirements of this study using data supplied on the JGI genome browser (https://genome.jgi.doe.go) for each gene accession (Data S1).

For investigating the transgenic PSR1-OE lines, FC were obtained for each line relative to UVM4 for d2, 3 and 6. Here, a list of biologically significant genes (OE-248) was obtained by including those where at least one time point was up or down by >2-fold for at least one transgenic line. In this case, each gene was designated either up or down according to which change had the greatest magnitude. Similar treatment was applied to the P-STRESS dataset (*16*) (time-points d3 and d5) so the two datasets could be compared by Venn diagram analysis (Data S1). In the latter case, P-adj values were not available hence the use of the FC cutoff. The OE-248 set was curated into functional roles and processes as above, annotating genes if necessary and considering *C. reinhardtii* biology (Data S1).

### Statistical analysis and Data processing

Statistical differences were evaluated by one-way ANOVA and by Tukey HSD test with a p-value of 0.05; both were performed using the software OriginPro (Version 2021, OriginLab Corporation, Northampton, MA, USA).

The growth rates and doubling times for each line were calculated according to (*52*). The specific growth rate (*μ = d^−1^)* was calculated for the exponential growth phase as follows: *μ=(ln(y_1_/y_0_))/(t_1_-t_0_)*, where y_1_ and y_0_ correspond to the biomass concentration values at the beginning and at the end of the exponential phase, respectively, and t_1_ and t_0_ are the days where y_1_ and y_0_ were obtained. The doubling time (d) was calculated as *ln(2)/μ*. Biomass productivity (*g L^-1^ d^-1^*) was determined with the equation *Bp= (Bc_f_ - Bc_i_/t*, where *Bc_i_*, and *Bc_f_* are the biomass concentration initial and final values for the cultivation time (t=7 d), respectively. The nutrient uptake and removal rates for each line were calculated according to (*53*). The maximum nutrient removal rates R_max_ (mg N L^-1^ d^-1^) were obtained by calculating the daily removal of a specific nutrient (CN_d_ - CN_d-1_) where CN_d_ is the nutrient concentration at a specific day and CN_d-1_ is the nutrient concentration on the day before, and finally selecting the highest removal rate observed on a specific day during the experiment. The nutrient consumption was calculated as *V (mg N g^-1^ dw) = (CN_0_-CN_1_)(Bc_1_-Bc_0_)*, where *CN_0_* and *CN_1_* are the media nutrient concentration values and *Bc_1_* and *Bc_0_* are the biomass concentrations at the early (t_*0*_=d_1_) and late exponential phase (t_*1*_ = d_4_). The nutrient uptake rate *k* (d^-1^) was obtained by dividing the nutrient consumption by the specific growth rate *μ*.

Multivariate analyses by PCA and Pearson’s correlation coefficient plots were carried out using PAST v4.08 (*54*). PCA was carried out on mean (n=3) values for log2(FC) values (Data S1) for two transgenic lines (8-27, 8-42) v. UVM4 control (6 data points per gene: 3 time points, 2 transgenic lines). PCA was also carried out on RPKM data (Data S1) for the above 3 lines, along with PI, PE and N data using data means (n=3) (9 data points per gene: 3 time points, 3 lines). Pearson’s correlation coefficients were also determined from these RPKM data in relation to the PI, PE and N data (Data S1).

## Acknowledgments

We would like to thank Ms. Carolina Lascelles, Ms. Morag Raynor and Dr. Ian Carr of the Next Generation Sequencing Facility (Leeds Institute of Biomedical & Clinical Sci.) for their hard work and advice. We are grateful Dr. Ruth Hughes and the Bio-imaging facility (University of Leeds) for helpful assistance. We are indebted to Dr. David Elliot for performing the ICMS analysis of ions in culture media. *Chlamydomonas reinhardtii* strain UVM4 was kindly provided by Prof. Dr. Ralph Bock.

## Funding

This work was supported by:

UK Research and Innovation (UKRI) BBSRC (BB/N016033/1).

ESRC funded GCRF Water Security and Sustainable Development Hub (ES/S008179/1) for the financial support provided to Miss Tatiana Zúñiga-Burgos.

The confocal microscopes in the Bioimaging Facility at the University of Leeds were funded by Welcome Trust grant WT104918MA.

## Author contributions

Project concept: AB, MACV

Transgenic constructs and transformation: LC, PM, MPD

Confocal microscopy: TZB, LC

Physiology experiments: TZB, SPS, LC

Western blotting, ICMS & assays: TZB

RNA-seq analyses & models: SPS

Investigation: TZB, SPS, LC

Supervision: AB, MACV, AGS

Figures: SPS, TZB

Writing—original draft: SPS, TZB

Writing-review & editing: SPS, AB, MPD, AGS, PM, TZB, MACV, LC

## Data and materials availability

All data are available in the main text or supplementary materials except for the RNAseq reads which are openly available from the University of Leeds Data Repository: https://doi.org/10.5518/1217

Supplementary Materials for

## This PDF file includes

Supplementary Text

Figs. S1 to S12

Tables S1 to S2

## Other Supplementary Materials for this manuscript include the following

Data S1

## Supplementary Text

### DNA construct for PSR1 over-expression (PSR1-OE)

The pLC8 DNA construct is shown in **Fig. S1** and the amino acid sequence of the fusion protein is shown in **Fig. S2**. The fusion protein consisted of *C. reinhardtii* PSR1 C-terminally fused to a 3xHA-tag followed by YFP and a Strep tag. The predicted MW was 109 kDa (1048 αα) and its expression was driven by a constitutive PSAD promoter (when under continuous light) and an RBCSi enhancer element. The construct was assembled into pUC57 backbone plasmid (kanR) using the synthesized PSR1 gene (Cre12.g495100.t1.1) including the 3xHA-tag. This plasmid was then used as a template for Golden Gate-based cloning which introduced PSADprom, RBSC2 intron, mVenus including the Strep tag and PSAD terminator. A second round of cloning introduced the ParoR gene components. The full cloning history is shown in **Fig. S3** and described in Materials and Methods in the manuscript. The pCL8 construct was then transformed into *C. reinhardtii* UVM4 by electroporation (Materials and Methods). Stable pLC8-transformed *Chlamydomonas* lines were screened for PSR1-YFP fusion mRNA by RT-PCR and by western blotting for levels of intact fusion protein (**Fig. S4**). The primers used for the construct assembly and screening are shown in **Tables S1, 2**.

### Nuclear targeting of PSR1-YFP fusion protein

Fluorescence confocal analysis was carried out on two of the PSR1-OE lines (8-27 and 8-42) to examine intracellular targeting of the PSR1-YFP fusion protein. The intracellular targeting and composition of the strongest PSR1-OE line 8-27 is shown in the manuscript (**Fig. 1**). Similar data for the weaker PSR1-OE line 8-42 is shown in **Fig. S5**. This demonstrated that there was nuclear expression of PSR1-YFP in both lines where the DAPI-DNA signal co-localized with PSR1-YFP in large organelles, consistent with the nucleus (**Fig. S5 B-D**). Unlike the nuclear DAPI-DNA signal, the DAPI-PolyP signal located close to the vacuole consistent with the known location of PolyP granules to the acidocalcisomes or vacuolar bodies (**Fig. S5 E-H**).

### PolyP accumulation is transiently increased with PSR1-OE

Comparison of PSR1-OE line 8-27 and the UVM4 control in a batch-culture time course indicated differences in the accumulation pattern of PolyP granules by fluorescence confocal microscopy (carried out as above). This was shown for single representative cells in the manuscript (**Fig. 1**). Here in **Fig. S6** images from multiple cells (10) are shown for the two lines for each time point. As indicated in the manuscript, PolyP granules were particularly large (<5 μm) with a very extensive but diffuse DAPI-PolyP signal apparent in the transgenic line at d2 compared with the control (**Fig. S6A**). However, by d3, granules had become smaller and more numerous in both lines but with more intense signal in the control compared with 8-27 (**Fig. S6B**). By d6, the signal was diffuse and weak in both lines (**Fig. S6C**).

### Growth rate and composition of medium and biomass during batch culture

Three PSR1-OE lines (8-2, 8-27, 8-42) were compared together with an untransformed UVM4 background control in a batch culture time course (standard conditions at 25°C under continuous light in nutrient-replete TAP medium: 1 mM P ~30 mg/L P). Growth in batch culture was compared in the manuscript in **Fig. 2A**, followed by depiction of PSR1-OE driven changes: medium P and Mg, and biomass P (**Fig. 2B-D**). In **Fig. S7**, measurements are shown for the parameters that did not change in transgenics versus control: growth rates (log plot); biomass chl a+b levels; pH; N (NH4^+^), S (SO4^2-^); Ca^2+^ and K^+^. A statistical analysis of the P in biomass differences between the transgenic PSR1-OE lines and the control is shown in **Fig. S8**.

### Functional analysis of gene expression changes during batch culture in the UVM4 control

RNA sequencing was performed on triplicate samples from d2, d3 and d6 for lines 8-27 and 8-42 and UVM4. Transcriptomic analysis focused first on the changes occurring in the background (strain UVM4) as it transitioned from log-phase (d3 v. d2) to stationary phase (d6 v. d2). Relative data was obtained (FC) using a 2-fold biological significance cut-off with a P-adj<0.05 statistical significance cut-off. This generated four sets of genes for early changes (d3 v. d2) and late (d6 v. d2) for both up and down (listed in **Data S1**). In the manuscript the data for the top 200 up or down genes were shown for the early and late comparisons (**Fig. 3A**). In **Fig. S9**, the full set of genes were analyzed in terms of function according to Protein Class Panther annotations available at JGI (*C. reinhardtii* genome v5.6 annotation file).

### Comparison of PSR1-OE data with a published P-stress experiment

The second focus of the transcriptional analysis was on the changes attributed to the PSR1-OE construct in the transgenics as compared to the UVM4 control during the batch-culture time course. This required a comparison of the transgenic lines (8-27, 8-42) against the UVM4 control. Fold-change (FC) gene expression data was determined at time points d2, 3 and 6. The greatest magnitude (i.e. the maxima or minima for each gene, whichever was greatest) were compared with the same from a published dataset from a P-starvation (P-STRESS) experiment (*16*). In the manuscript a biological significance cutoff of 2-fold for both datasets was used. This cutoff generated a subset of 248 genes (OE-248) for our data (98% p-adj<0.1; 68%, p-adj<0.05) (**Data S1**) but gave a much larger subset of ~4000 genes with the P-STRESS data (PS-4354). In the manuscript, good agreement was seen between the OE-248 and P-STRESS data sets (60% of the OE-248 set were also altered in the P-STRESS set). Here in Supplemental, the same exercise was repeated using similar numbers of genes from both data sets, to confirm the findings. The OE-248 data set (>2-fold cutoff) had 177 up genes and 71 down genes. To compile a list of similar numbers from the P-STRESS dataset, modified integer cut-offs of 6-fold (up) and 25-fold (down) were applied to the P-STRESS dataset with the results shown in **Fig. S10**. For the 177 up genes of the OE-248 set, 25% showed agreement with the P-STRESS data with a >6-fold cut-off (cf. 68%>2-fold). With the strongest line 8-27 the agreement for up genes was now 30% compared with 70% (>6-fold cutoff cf. >2-fold cutoff of P-STRESS data) whereas with line 8-42 it was 19% compared with 51%. This indicated a continued agreement with the higher ranked up genes from the P-STRESS dataset. In contrast, with the down genes, only three genes (FDX5, XBAT33 and RBCS2) from the OE-248 dataset (4%) showed agreement with a similar number of down-regulated genes from the P-STRESS data (c.f. 46% with a >2-fold cut-off). This reinforced the findings in the manuscript that PSR1-OE mimicked more closely P-stress for the upregulated cohort of genes rather than the down-regulated genes.

**Fig. S1.**
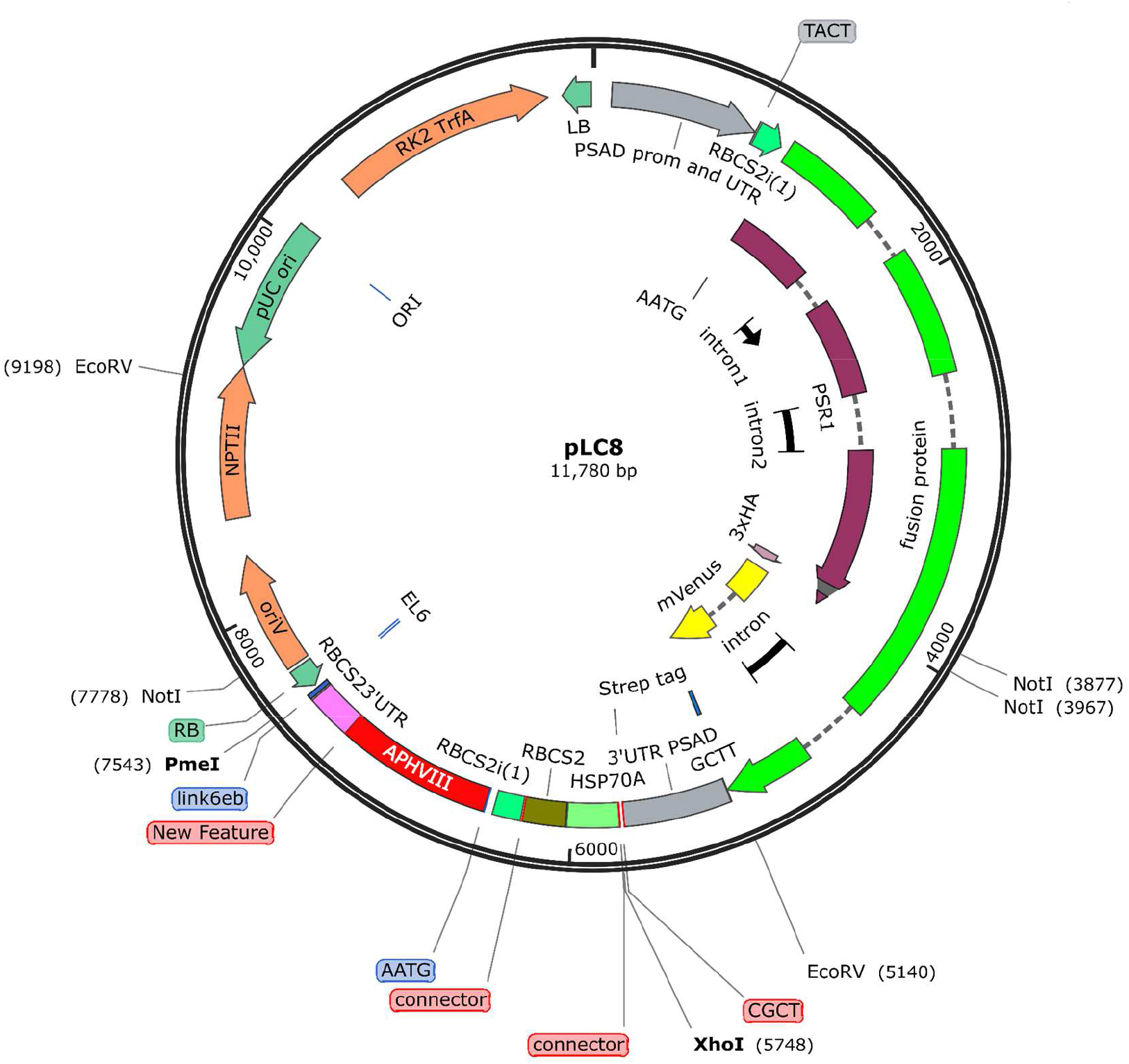
Plasmid map of pCL8, the PSR1-OE construct. The pLC8 construct was based on the pUC57 backbone, which provided kanamycin resistance (nptII) for microbial selection. From the top, the PSR1-OE component included the PSAD constitutive promoter with an RBCS2i enhancer driving the PSR1 gene. The latter was terminally fused to a 3X HA tag then YFP (mVenus) followed by 3’UTR of PSAD. Downstream of this gene was a second gene construct for Paromycin resistance for enabling selection of algal transformants.

**Fig. S2.**
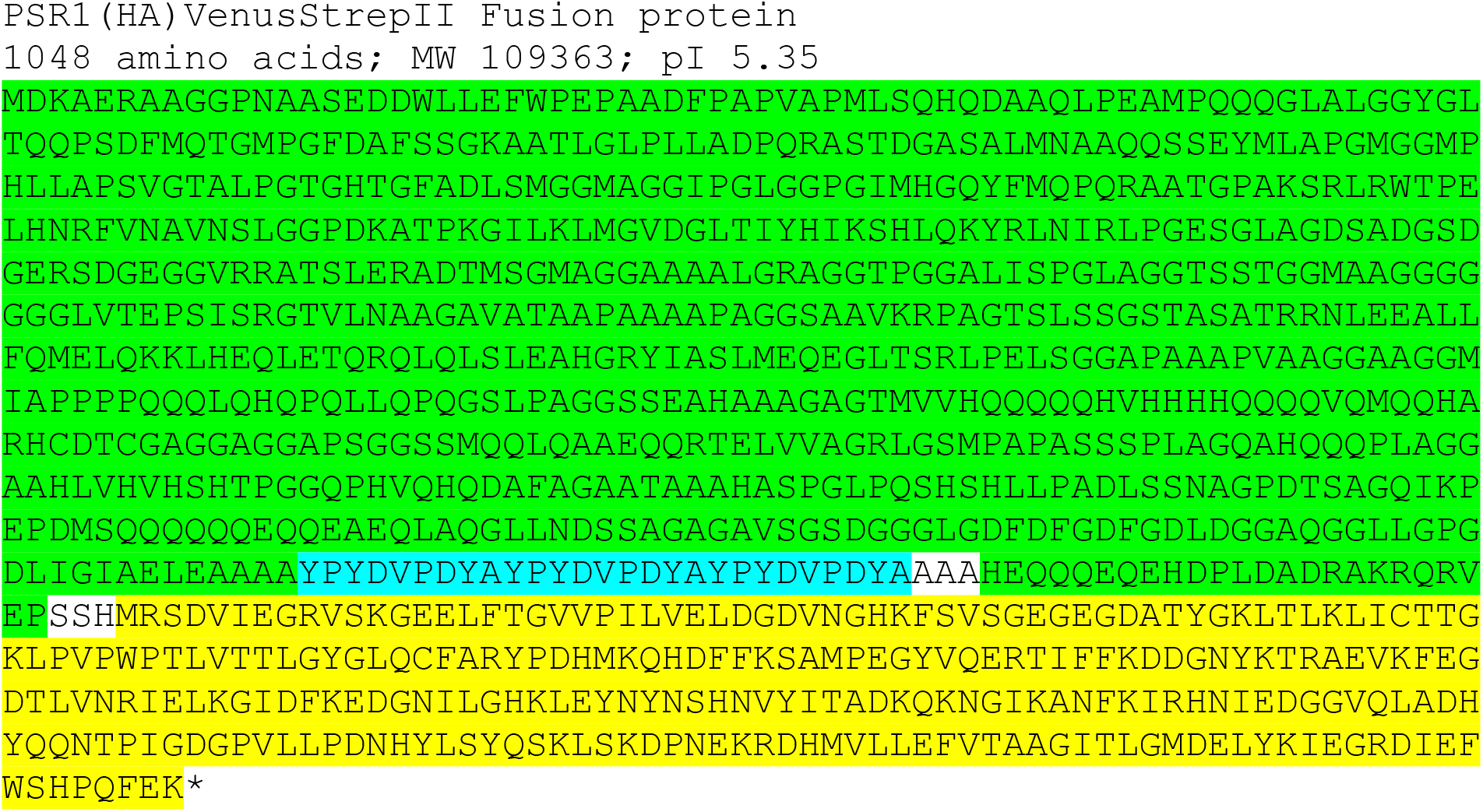
Amino acid sequence of PSR1-OE construct fusion protein. The fusion protein was of predicted size 109 kDa and consisted of an N-terminal fusion of PSR1 (MDK…VEP) (*C. reinhardtii* v5.6 Accession: Cre12.g495100.t1.2) containing a 3xHA tag inserted towards the end in a stretch of Alanines (YPY…DYA), with a C-terminal mVenus YFP (MRS…FEK).

**Fig. S3.**
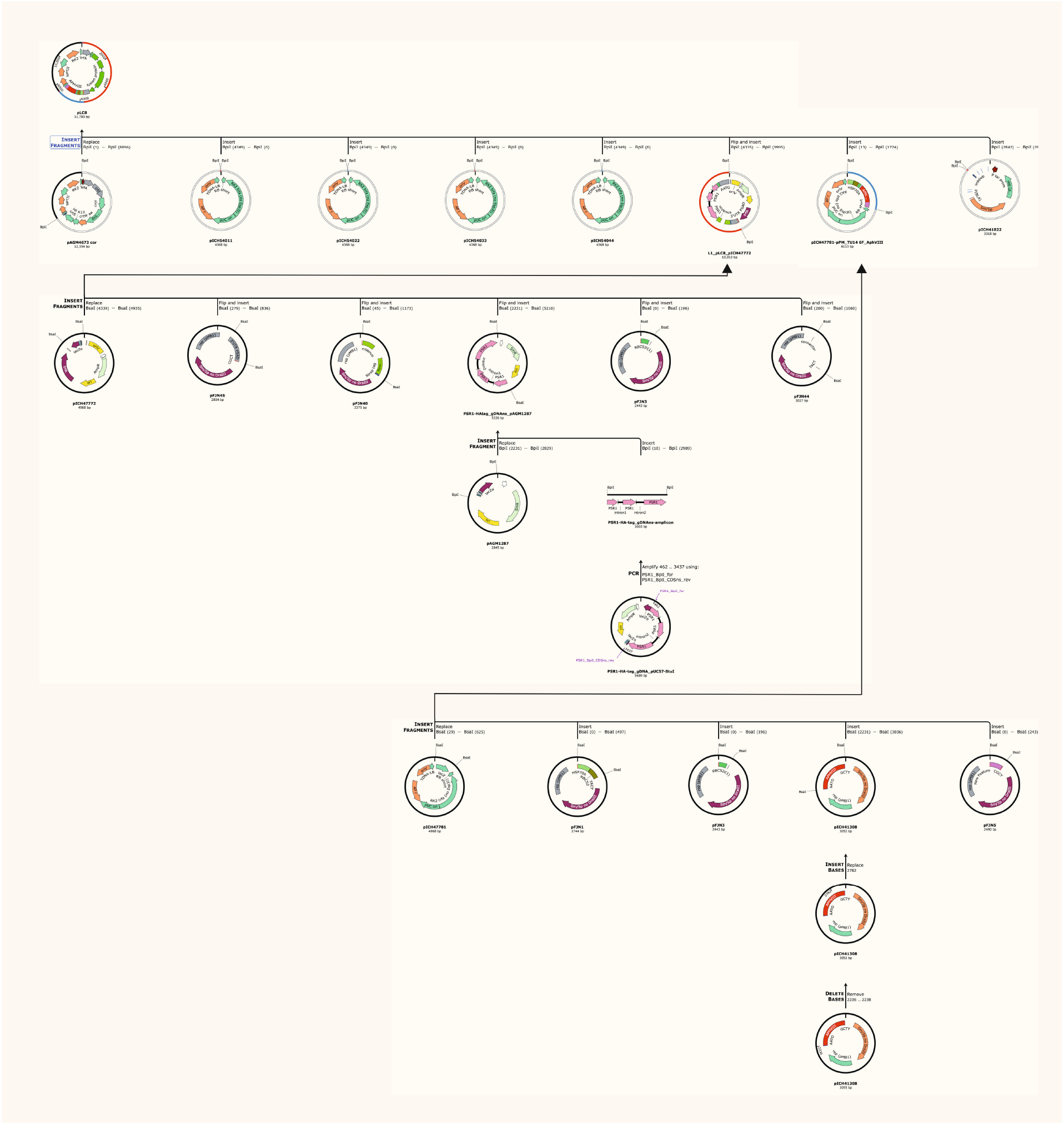
Full cloning history of the pCL8 construct (zoom into image). The PSR1 gene (Cre12.g495100.t1.1) with an inserted 3xHA-tag was synthesized (Genscript Biotech Corporation, UK) and cloned into pUC57 via the StuI restriction site. This plasmid was then used as a template for Golden Gate-based cloning. The following level 0 plasmids were used: pCM0-001 (PSAD prom), pCM0-024 (RBCS2 intron), pCM0-044 (mVenus, incl. Strep-tag), pCM0-114 (PSAD term) all from (*47*), and L0_PSR1 (PSR1 cloned into pAGM1287 (MoClo Plant Kit) in this study). For the generation of level 2 plasmids for Chlamydomonas transformation, the following level 1 plasmids were used: pAGM4673 (L2 backbone, MoClo Plant Kit), pICH41822 (L2 end-linkers MoClo Plant Kit), pICH54011, pICH54022, pICH54033, pICH54044 (Dummies, MoClo Plant Kit), pCM1-27 (ParoR,) (*47*) and L1_PSR1 (PSADprom-RBCS2intr-PSR1-mVenus-PSADterm, this study).

**Fig. S4.**
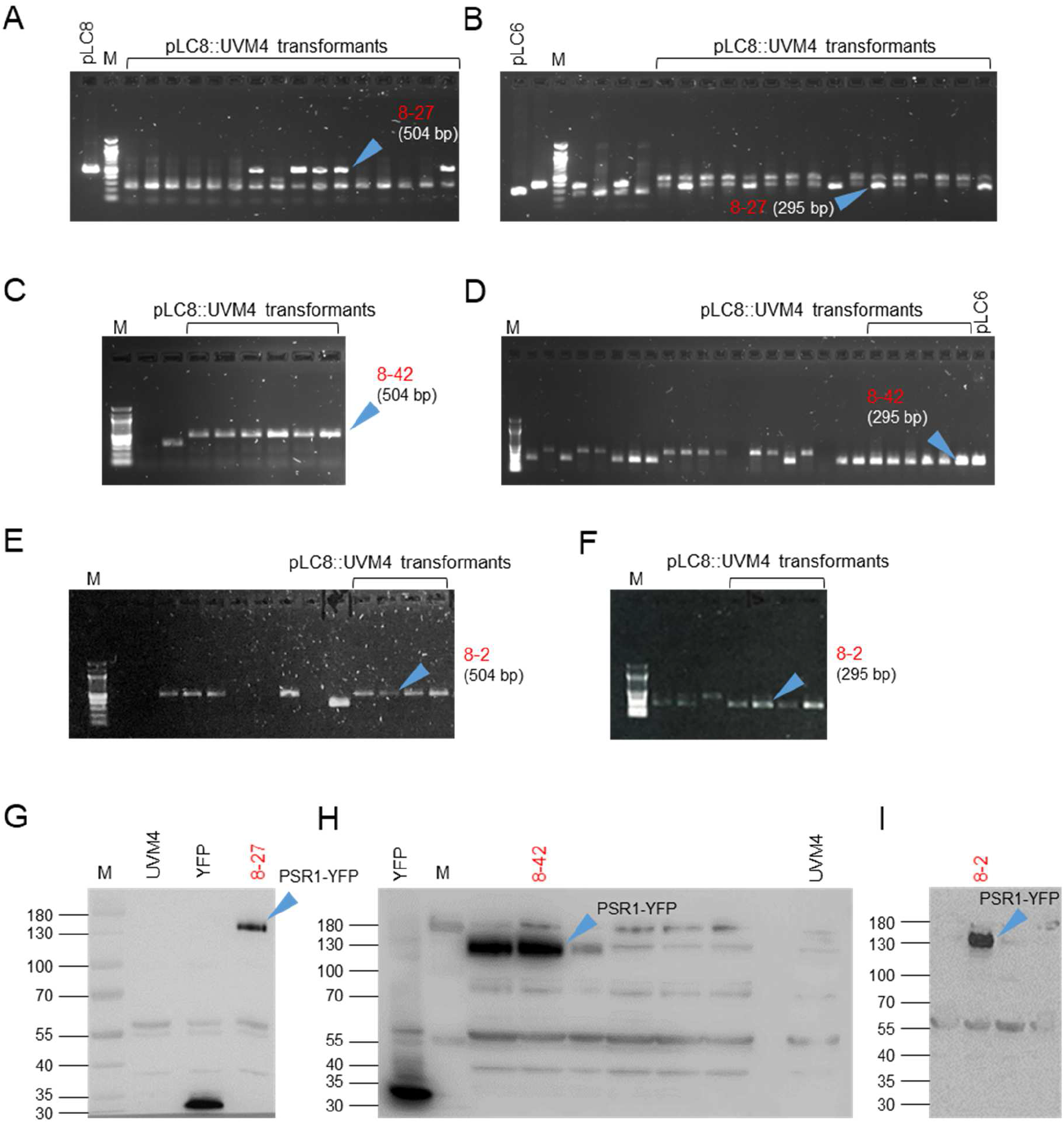
Screening for PSR1-YFP construct and fusion protein in transgenic lines. Colonies from three independent transformations of *C. reinhardtii* were screened by (**A-F**) colony PCR for the presence of the PSR1-YFP construct and (**G-I**) Western blotting for presence of the PSR1-YFP fusion protein. (**A, B, G**) **Line 8-27** is highlighted from transformation date 23-11-2017. (**C, D, H**) **Line 8-42** is highlighted from transformation date 22-03-2018. (**E, F, I**) **Line 8-2** is highlighted from transformation date 26-04-2018. (**A, C, E**) The PSR1 portion of the construct was detected by PCR primers LC40 and LC42 giving rise to a 504 bp product, utilizing the pLC8 plasmid DNA as a positive control. (**B, D, F**) The Venus YFP portion of the construct was detected using primers LC43 and LC45 generating a 295 bp product, using pLC6 plasmid DNA as a positive control. (**G-I**) Western blot analysis of protein extracts from the selected colonies for presence of the fusion protein using anti-YFP antibody (predicted size 109.4 kDa). The controls were untransformed UVM4 (negative) and UVM4 transformed with the YFP construct only (positive). Chlorophyll loadings were 2.5 μg (**G, I**) and 5.0 μg (**H**). Marker (M), primers are listed in **Table S1**.

**Fig. S5.**
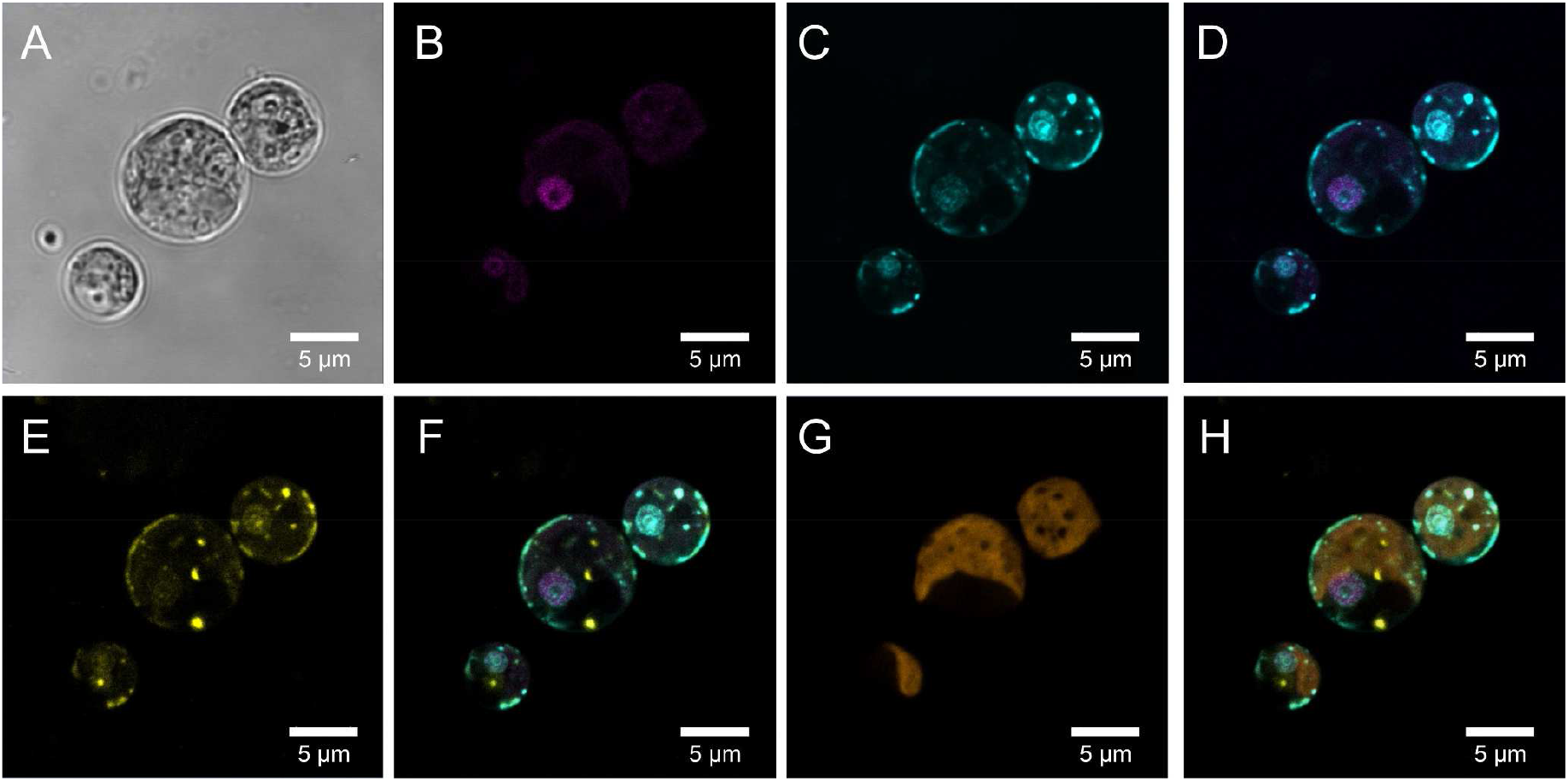
Intracellular targeting of PSR1-YFP fusion protein as determined by fluorescence confocal microscopy. Intracellular location of the PSR1-YFP fusion protein shown in (**A-H**) for representative cells from PSR1-OE line 8-42 grown in TAP media. Similar data for the expression of the independently transformed PSR1-OE line 8-27 is shown in **Fig. 1**. (**A**) bright-field images indicating cell diameter. (**B**) Venus-YFP signal (Emission λ 520-550 nm: magenta) indicating location to the nucleus which was identified by DAPI-DNA fluorescence (Emission λ 420-475 nm: cyan) (**C**) followed by co-localization of the DAPI and YFP signals in the merged image (**D**). The PolyPhosphate (PolyP) granules are indicated by the DAPI-PolyP (Emission λ 535-575nm: yellow) (**E**). These are shown to be separate entities from those identified by DNA-DAPI staining in the merged image (**F**). Chlorophyll UV-fluorescence (Emission λ670-720 nm: orange) indicating the single large cup-shaped chloroplast (**G**) and the merged image (**H**) which excludes the YFP and PolyP signals from this organelle, placing the PolyP signal to the periphery of the dark central region of the cell (vacuole).

**Fig. S6.**
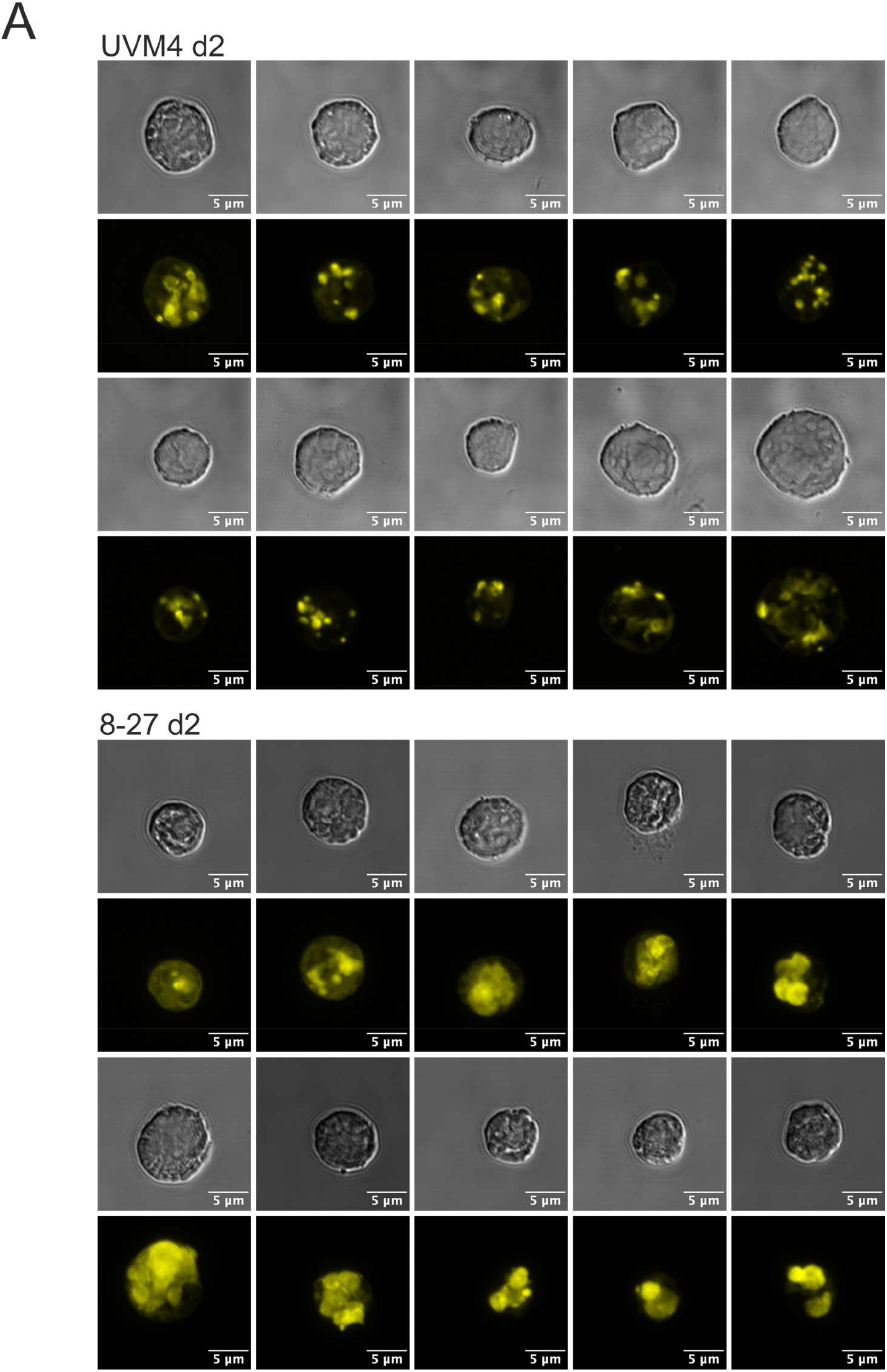
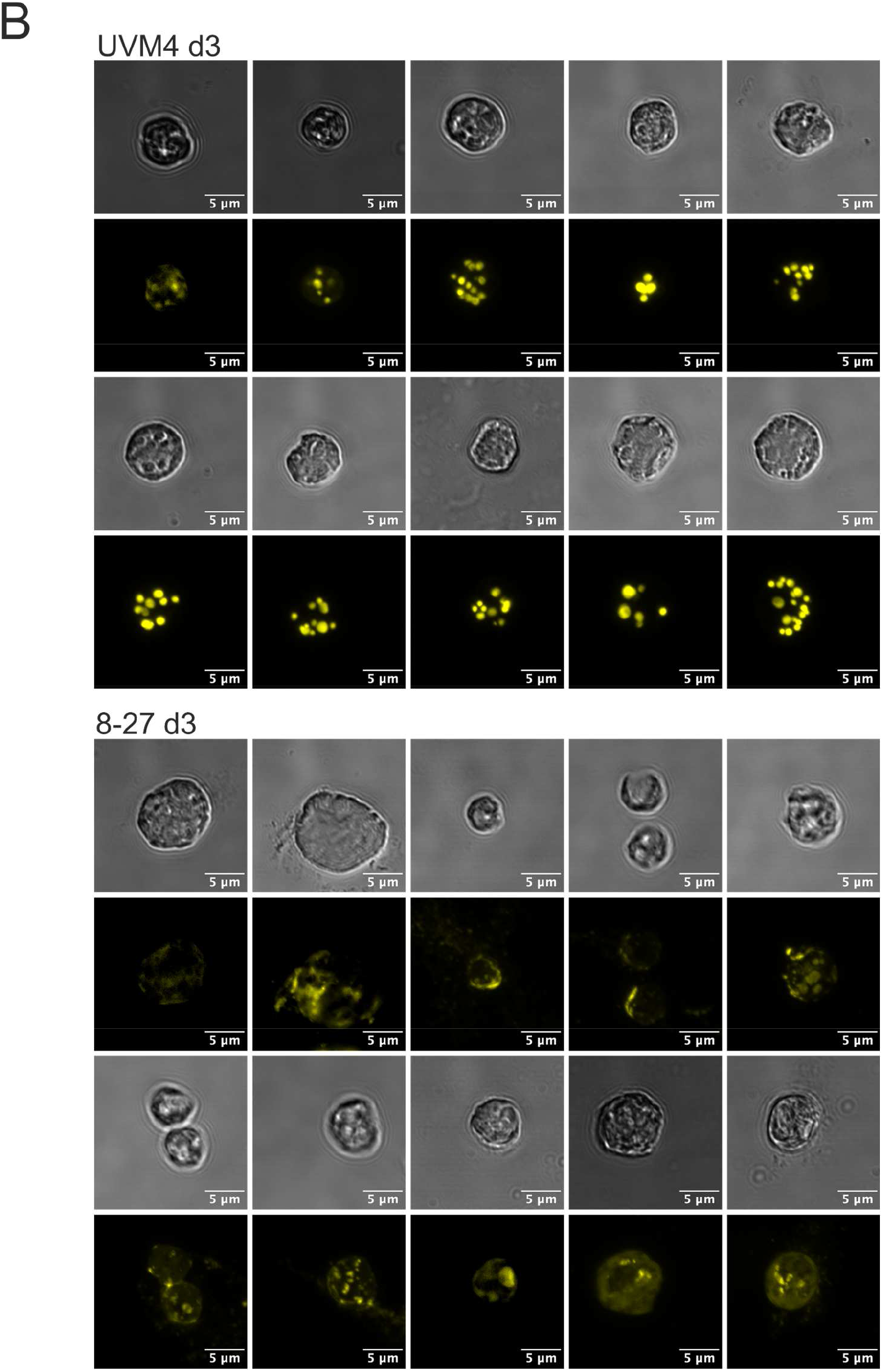
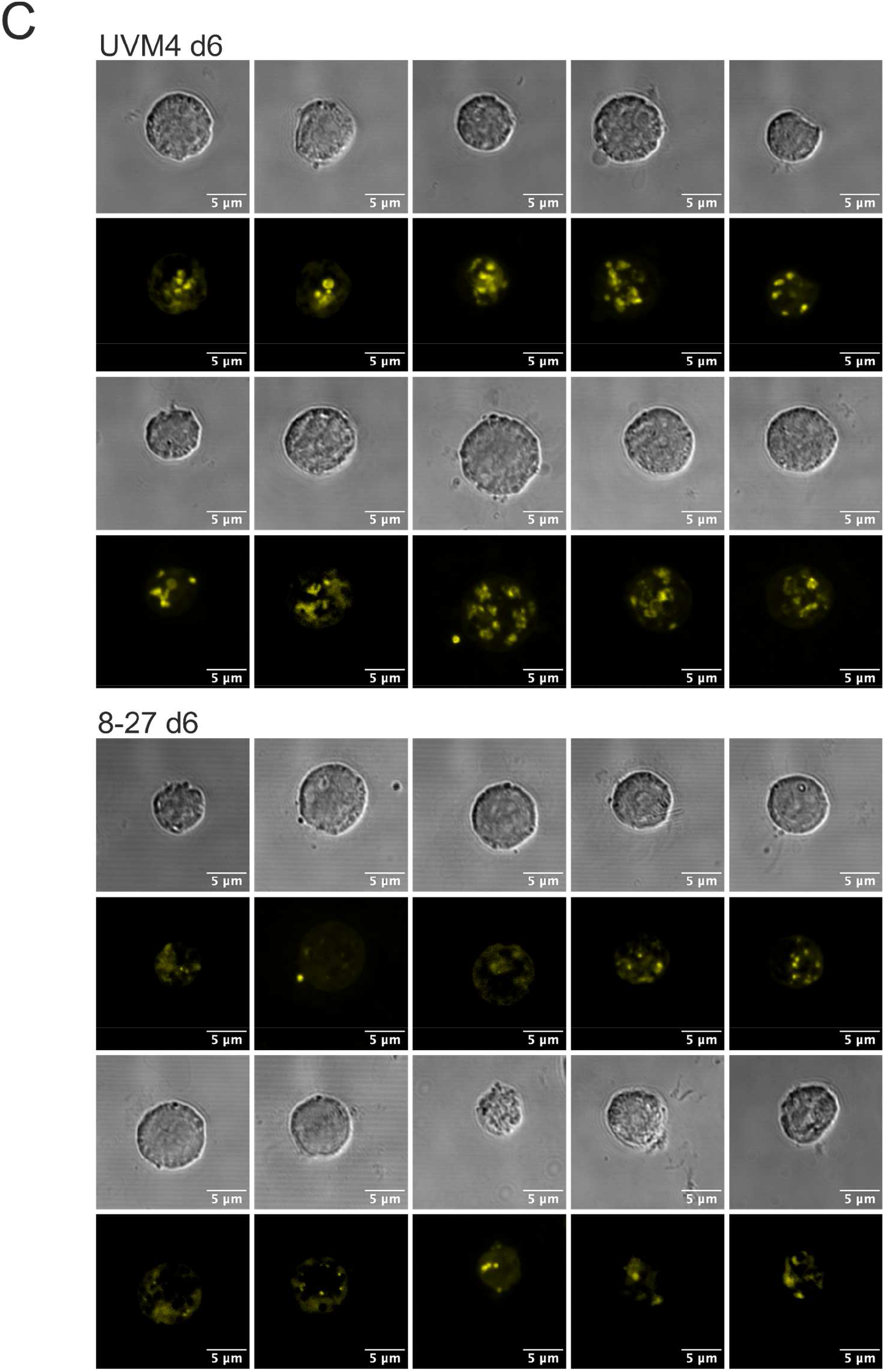
Increases in PolyP storage granule size as determined by fluorescence confocal microscopy associated with PSR1-OE. Increases in PolyP storage granule size as determined by fluorescence confocal microscopy associated with PSR1-OE. Displayed in (**A-C**) are differences in the accumulation of PolyP in cells from a batch culture time course in TAP media for time points d2 (**A**), d3 (**B**) and d6 (**C**). At each time point 10 representative cells are compared for control line UVM4 (top) and PSR1-OE line 8-27 (bottom). For each cell the bright field (top) and the DAPI-PolyP signal (bottom) is shown (Emission λ 535-575nm).

**Fig. S7.**
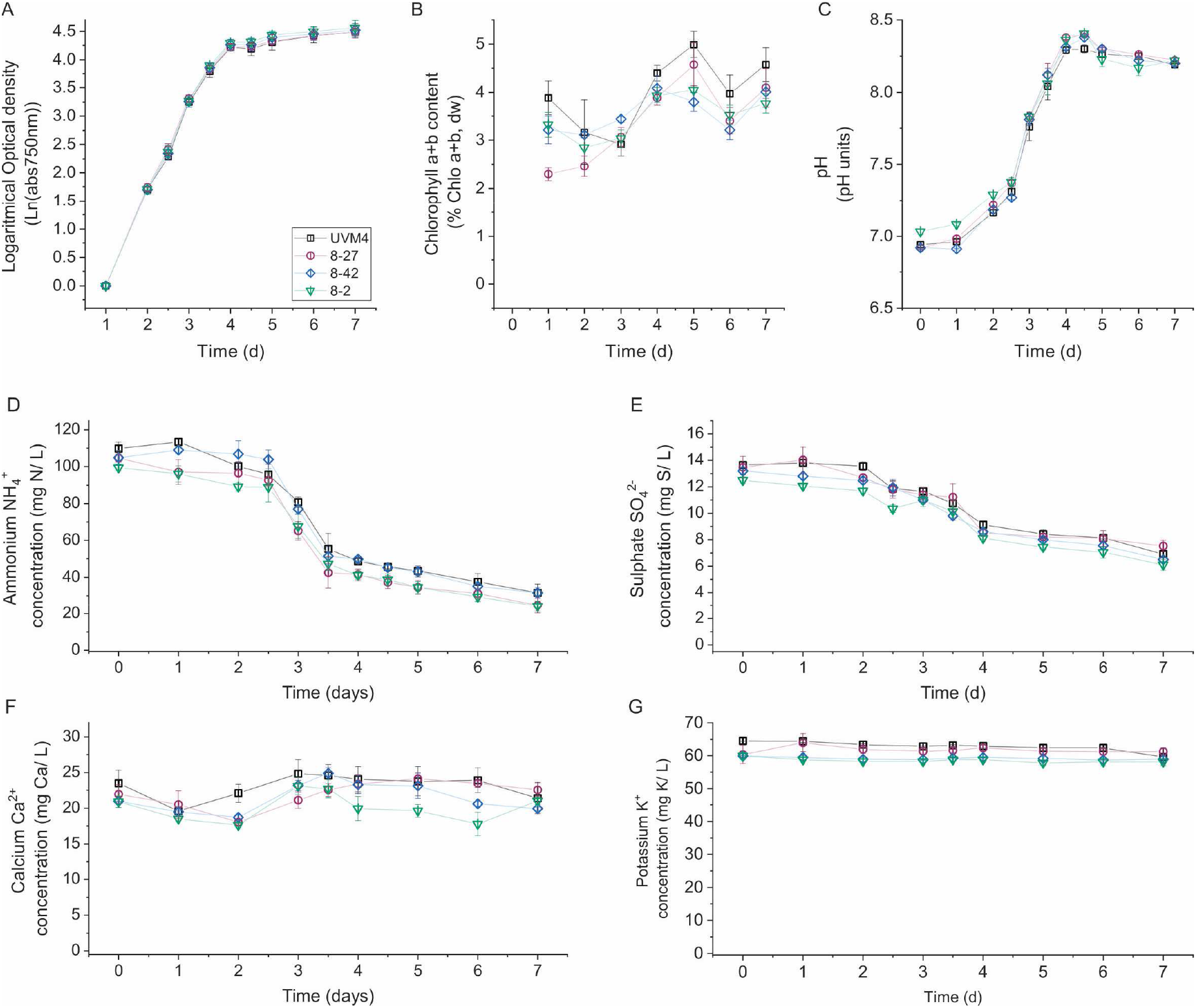
Physiological parameters of transgenic lines and UVM4 control. Three transgenic PSR1-OE-lines, along with untransformed UVM4 background control, were cultivated under small-scale batch culture conditions in TAP media under continuous light. (**A**) Growth rates (log plot of optical density (OD)), (**B**) biomass chl a+b levels, (**C**) pH, (**D**) Medium N levels, (**E**) medium S levels, (**F**) medium Ca levels and (**G**) medium K levels.

**Fig. S8.**
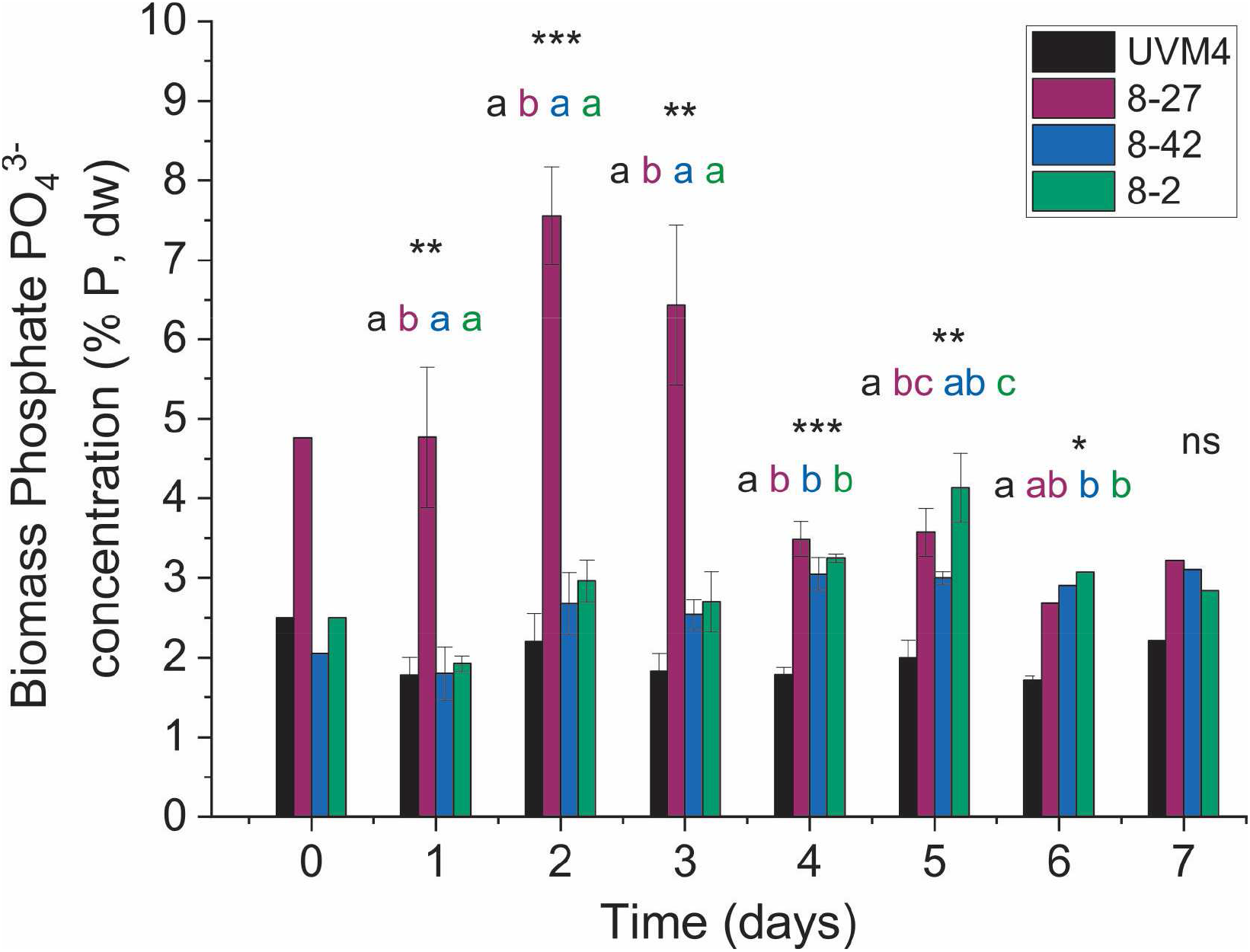
Statistical analysis of enhanced P in biomass accumulation in PSR1-OE lines. Measurements P in biomass composition at different stages of algal growth. Three transgenic PSR1-OE-lines, along with the untransformed UVM4 background control, were cultivated under small-scale batch culture conditions in TAP media (30 mg/L P ~ 1mM) in continuous light. Statistical differences were analysed by One-way ANOVA to validate the replicates and determine differences between the transgenic lines and the control: ns (not significant), * (P<0.05), ** (P<0.01), *** (P<0.001) and **** (P<0.0001). Where p<0.05 (One-way ANOVA), multi-comparison statistical analysis was carried out between each of the four lines with a Tukey HSD test (p-value<0.05). Letters from a-d were assigned to group the lines with no significant differences on each day of cultivation.

**Fig. S9.**
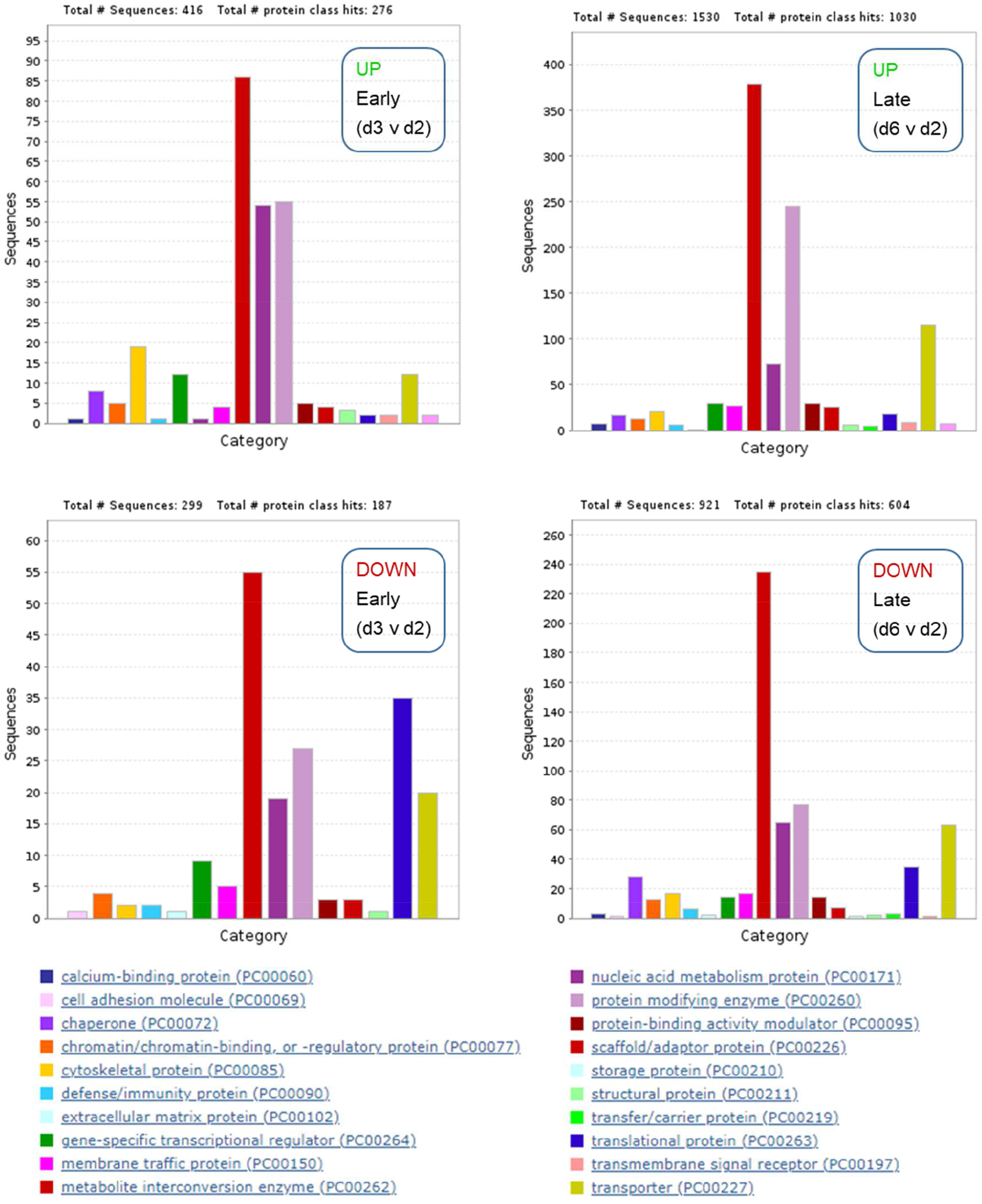
Gene function analysis of UVM4 during batch culture. Analysis based on Panther gene ontology for Protein Class for early and late, up and down as labelled. Relative data was obtained (FC) using a 2-fold biological significance cut-off with a P-adj<0.05 statistical significance cut-off. This generated four sets of genes for early changes (d3 v. d2) and late (d6 v. d2) for both up and down (listed in **Data S1**). Protein Classes are indicated in the legend and numbers falling into these classes in relation to the total is indicated at the top of each chart. Majority changes at all stages, up or down were Metabolite-interconversion enzyme genes, followed by Protein modifying enzyme genes, Nucleic acid metabolism proteins and Transporters. Large early decrease are shown in Translational protein and Transporters, and a relatively large increase is shown in Cytoskeletal protein genes.

**Fig. S10.**
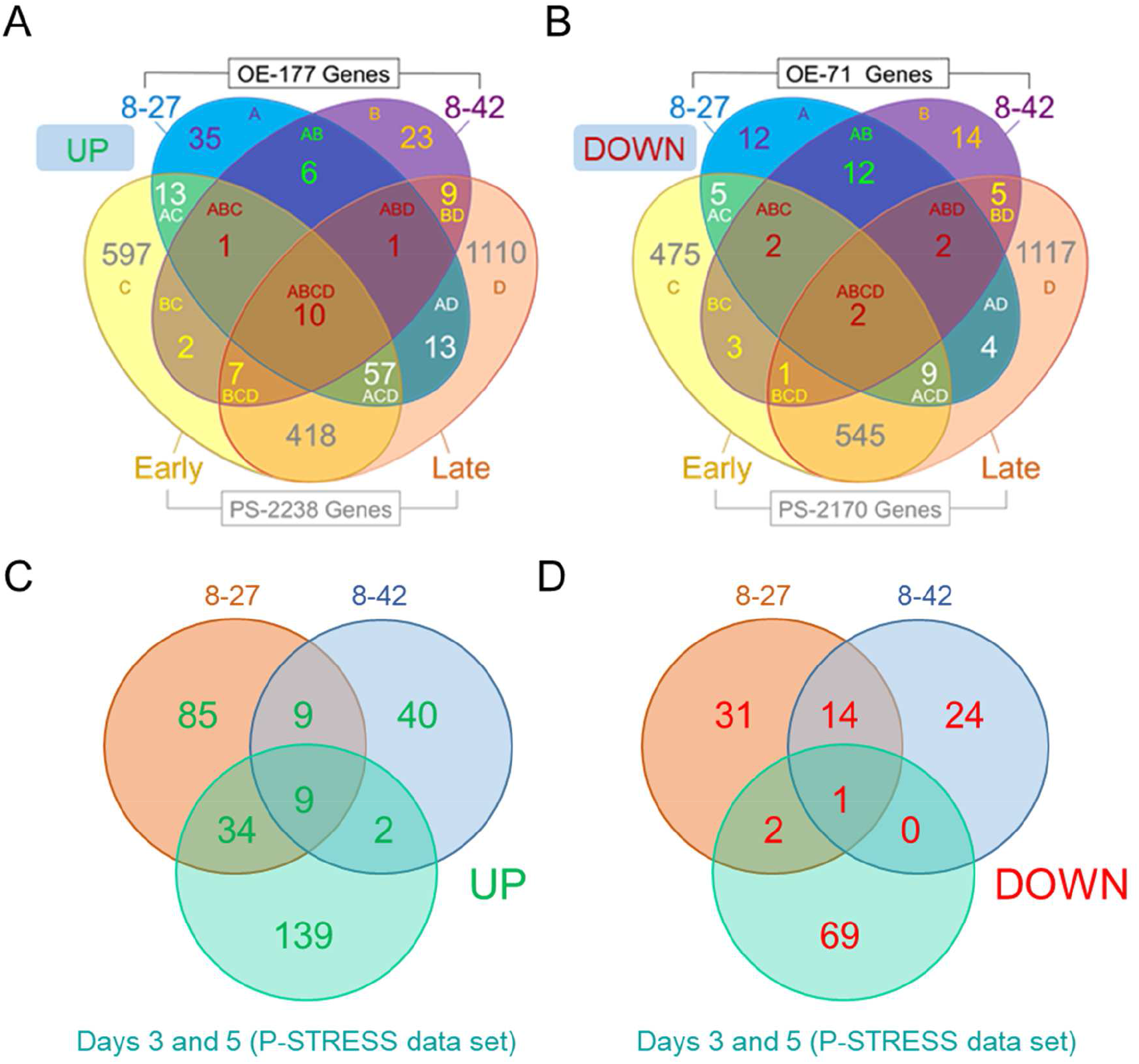
Co-regulated expression changes between two PSR1-OE lines (8-27 and 8-42) and the P-STRESS data showing direction of regulation (up or down). Co-regulation occurrences are shown for maximum (**A, C**) or minimum (**B, D**) FC values (≥2-fold FC relative to UVM4) for a given gene from the PSR1-OE lines in comparison with the P-STRESS dataset FC’s. (**A, B**) Gene number per Venn sector is indicated and each sector is labelled as follows: A (8-27), B (8-42), C (P-STRESS d3 “EARLY”) and D (P-STRESS d5 “LATE”). Here the P-STRESS FC cut-off was also ≥2-fold. (**C, D**) Similar analysis employing different cut-offs for the P-STRESS dataset FC’s (**C**) ≥ 6-fold and (**D**) ≥ 25-fold to generate equivalent gene numbers as observed for OE-248 up (177) and down (71).

**Fig. S11.**
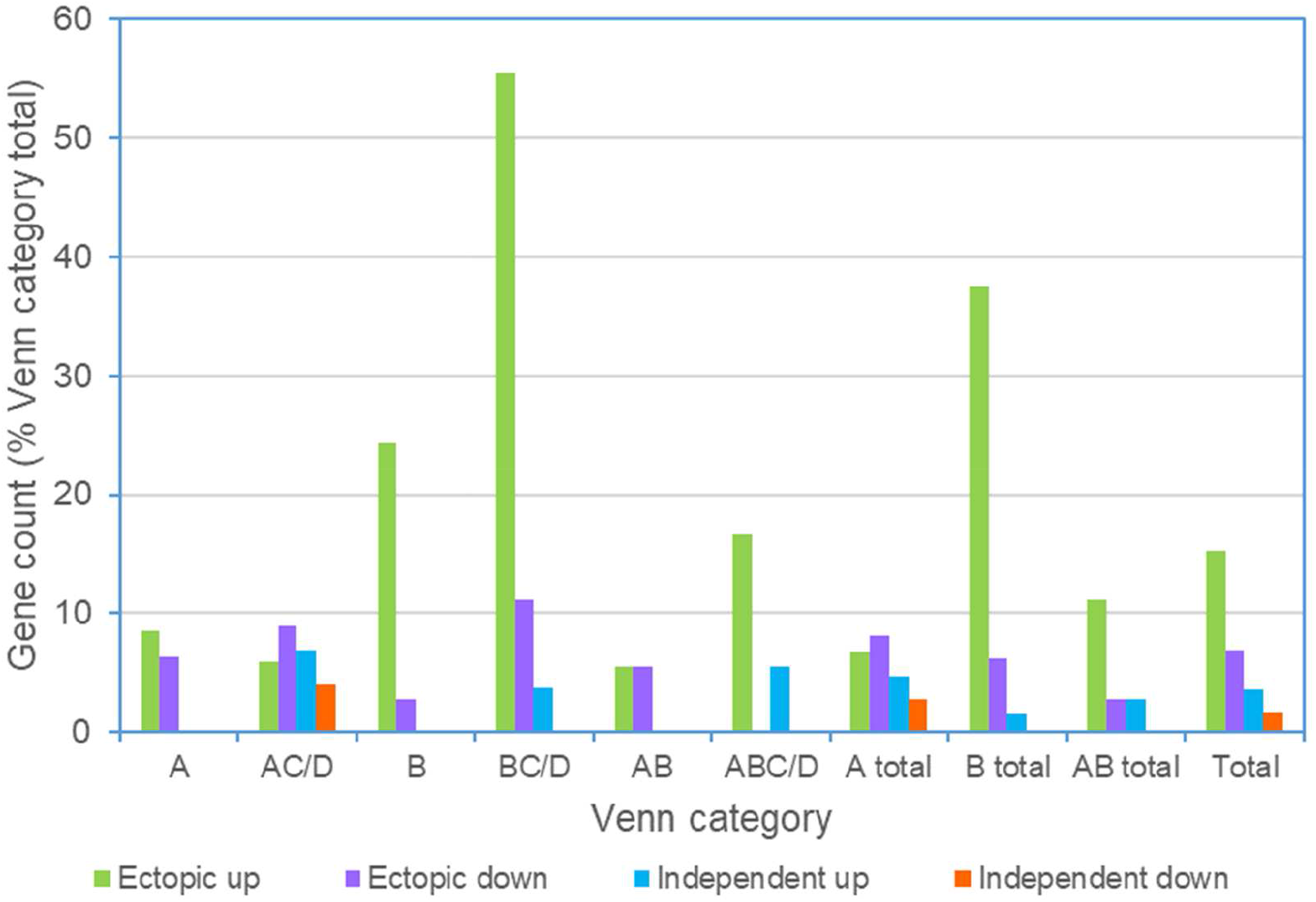
Frequency of ectopic gene regulation in the *psr1-1* mutant among OE-248 genes. The data refers to the behaviour of the OE-248 gene list in the *psr1-1* mutant. These data were published as part of the P-STRESS dataset where P-stress was not just carried out on a strain with wild-type PSR1 (Venn C/D) but also provided for the *psr1-1* mutant (FC: P-stress v. no stress) (**Data S1**) (*16*). Note that up or down refers to behaviour in the *psr1-1* mutant under P-stress in the P-STRESS dataset. Gene counts are shown for each Venn category which was defined by the PSR1-OE FC values (v. UVM4) where A refers to line 8-27 and B to line 8-42. Here, Venn subsets refer to a comparison of these lines within OE-248 (A, B) and the published P-STRESS data set (C/D) (**Fig. 3B**). In **Data S1**, mutant and wild-type P-STRESS data were compared for each of the OE-248 genes. Note that genes in Venn sets A, B, AB were not significantly changed in gene expression in the P-STRESS data set (FC<2 and >0.5) for the wild-type PSR1 strain (C/D), so where these genes were significantly up/down regulated in the mutant, this was considered ectopic; the remainder counts being P-stress unregulated genes (not shown). In Venn sets XC/D, genes were defined as PSR1-independent where FC values were (i) similar (ΔFC <2-fold) in wild-type and mutant (ii) in the same direction (up or down) and (iii) biologically significant in both cases (FC>2 or <0.5) in the P-STRESS dataset. Genes in Venn XC/D were defined as ectopic where the FC’s were 2-fold greater in the mutant relative to wild-type (same or opposite direction). Here the remainder counts were PSR1-dependent P-stress regulated (FC>2 and <0.5) (not shown).

**Fig. S12.**
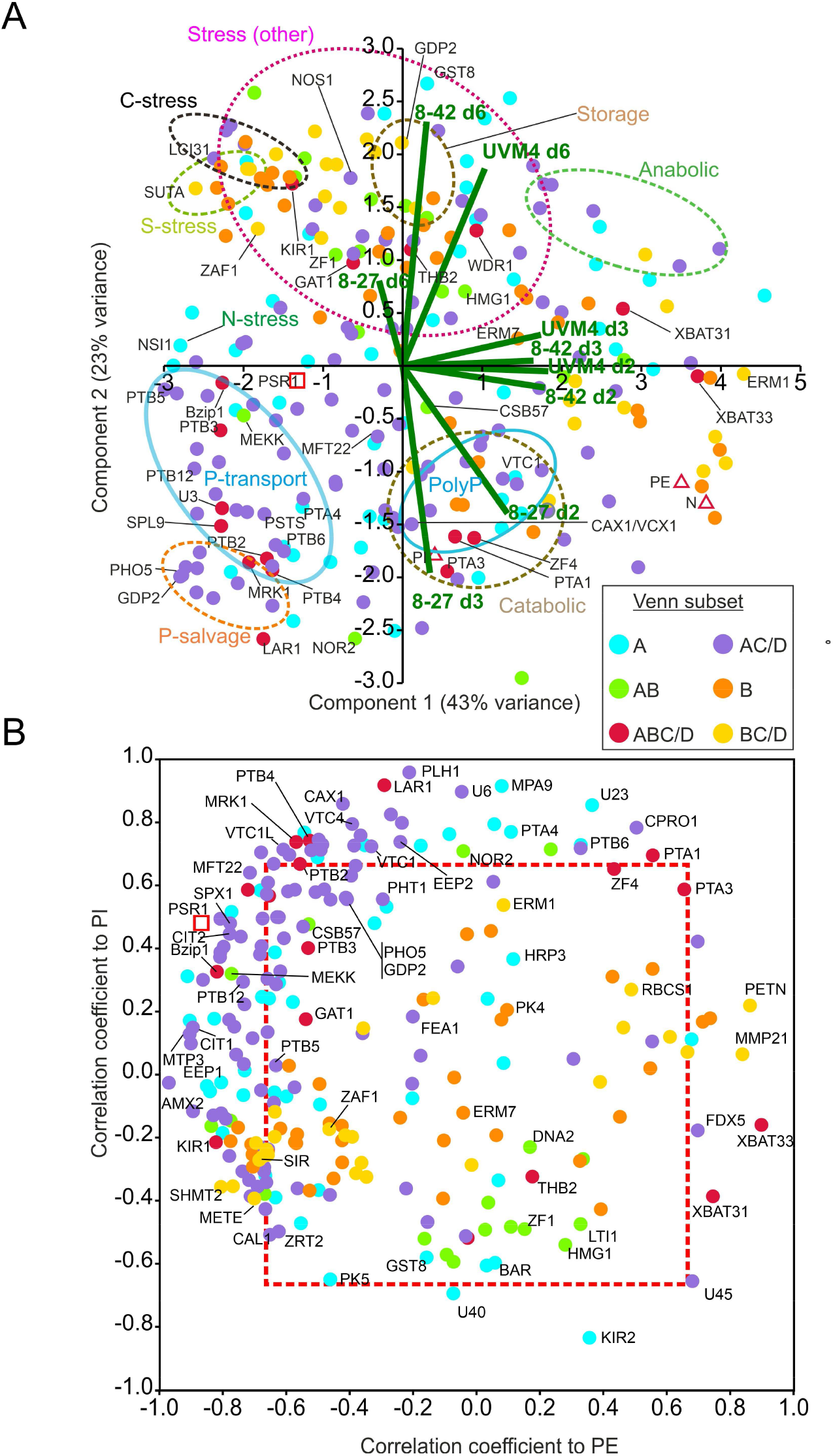
Temporal regulation of PSR1-OE gene expression. Patterns of gene expression focusing on temporal factors are shown for the OE-248 gene set. The charts are coded for the six Venn diagram sector subsets (inset) as described in **Fig. 3B**. The data point for the full PSR1 gene mRNA (not specific to endogenous gene or transgene) is shown (□). (**A**) PCA analysis of normalized mean (n=3) RPKM and PE, PI and N measurements (Δ). Biplots (-) shown for three lines and timepoints. Clusters highlighting functional processes are encircled. (**B**) Plot of Pearson’s correlation coefficients for RPKM data v. PI and v. PE for each OE-248 gene. Coefficients outside the boxed region (---) were significant for PI or PE (P<0.05). In all cases mean data was from n=3 culture replicates.

**Table S1.**
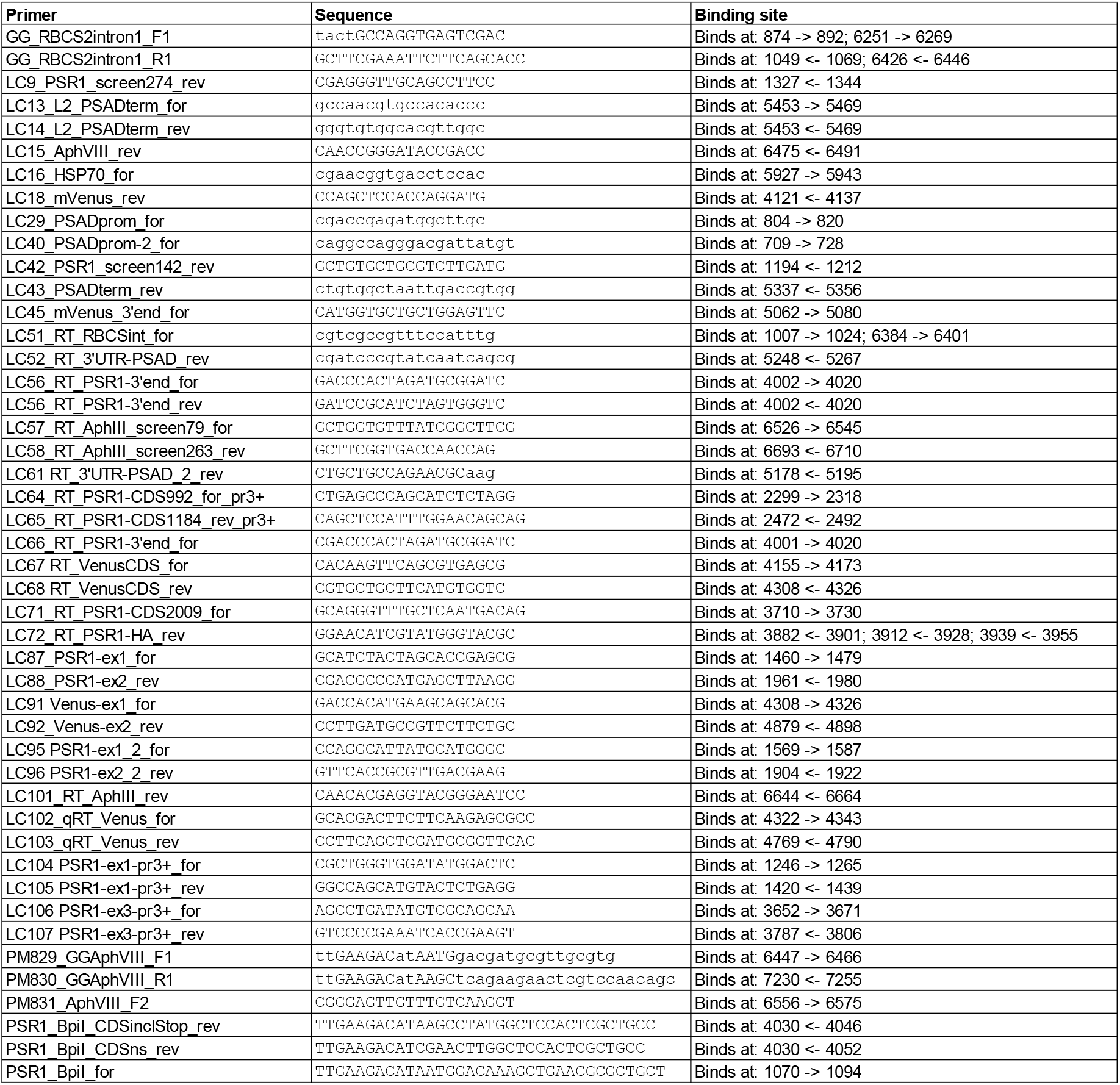
Primers used in cloning and screening of transformants.

**Table S2.**
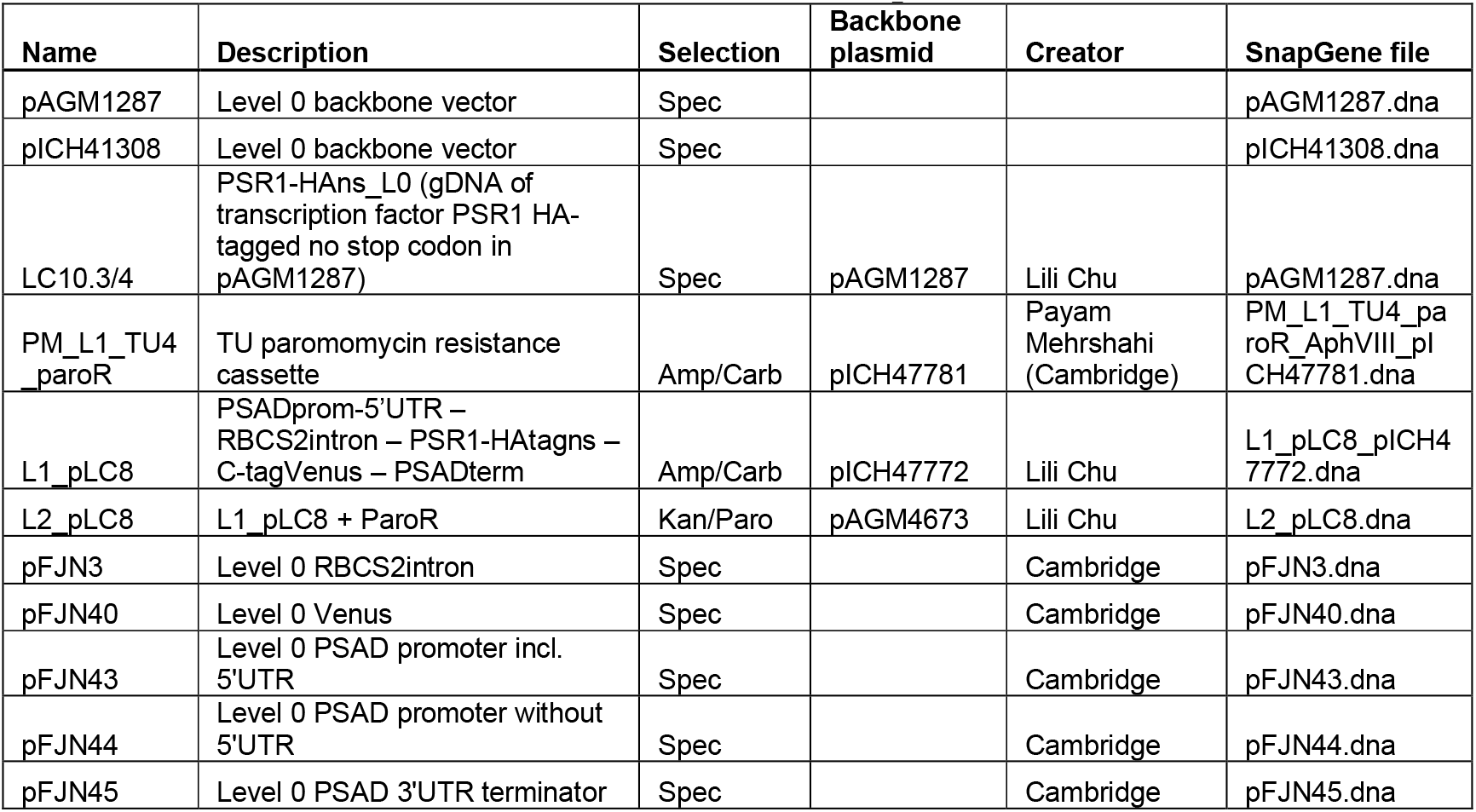
Plasmids used in the construction of the pLC8 construct.

